# SOX2 is required independently in both stem and differentiated cells for pituitary tumorigenesis in *p27* null mice

**DOI:** 10.1101/2020.06.09.142638

**Authors:** Veronica Moncho-Amor, Probir Chakravarty, Christophe Galichet, Ander Matheu, Robin Lovell-Badge, Karine Rizzoti

## Abstract

Loss of P27 predominantly results in development of murine pituitary intermediate lobe (IL) tumours. We previously showed that the pleiotropic protein P27 can drive repression of the transcription factor *Sox2*. This interaction plays an important role during development of *p27^-/-^* IL tumours because loss of one copy of *Sox2* diminishes tumorigenesis. Here, we have explored the cellular origin and mechanisms underlying melanotroph tumorigenesis in *p27^-/-^* IL. We show that IL hyperplasia is associated with reduced cellular differentiation, while levels of SOX2 increase in both stem cells (SC) and melanotrophs. Using loss-of-function and lineage tracing approaches, we demonstrate that SOX2 is required cell-autonomously in *p27^-/-^* melanotrophs and SCs for tumorigenesis. This is supported by studies deleting the *Sox2* regulatory region 2 (*Srr2*), which is the target of P27 repressive action. Single cell transcriptomic analysis reveals that activation of a SOX2-dependent MAPK pathway in SCs is important for *p27^-/-^* tumorigenesis. Our data highlight different roles of SOX2 following loss of *p27*, according to the cellular context. Furthermore, we uncover a tumor-promoting function for SCs, which is SOX2-dependant. In conclusion, our results imply that targeting SCs, in addition to tumour cells themselves, may represent an efficient anti-tumoral strategy in certain contexts.

## Introduction

Tumour formation involves a complex interplay of cellular interactions. Most frequently, transformed cells interact with their microenvironment, which provides a supportive niche for the formation, maintenance and dissemination of the tumour (Baghban et al., 2020). In addition, the tumours often display considerable cellular heterogeneity, which can include the presence of cancer stem cells (CSCs). These are a subpopulation of undifferentiated tumoral cells that may originate from resident tissue stem cells (SCs), and which fuel growth and recurrence of the tumour (Lytle et al., 2018). In rarer cases, cells acquiring mutations may not form the tumour themselves, but instead induce others to do so. This has been observed in the murine intestine (Balbinot et al., 2018), liver (Lujambio et al., 2013), skin (Mescher et al., 2017), in leukaemia (Kode et al., 2014), and in the pituitary where tissue SCs harbouring a Wnt pathway activating mutation recruit surrounding cells to form tumours resembling human craniopharyngiomas (Andoniadou et al., 2013).

The pituitary is a small endocrine gland located just under the brain, connected to the hypothalamus that controls its secretions. In the mouse it comprises three lobes, the posterior lobe, comprising axon terminals and glial cells, the anterior lobe (AL), which is highly vascularized and contains most endocrine cell populations, and the intermediate lobe (IL), which is avascular (Tanaka et al., 2013) and populated by a single endocrine population; these are melanotrophs that secrete melanocyte stimulating hormone (MSH). MSH is processed from its precursor pro-opio-melanocortin (POMC), which is also produced in AL by corticotrophs, where it is cleaved into adrenocorticotropic hormone (ACTH) (Drouin, 2016). Expression of *Pomc* is controlled in both cell types by the T-box factor TBX19 (Lamolet et al., 2001). In the embryo, the pioneer transcription factor PAX7 controls acquisition of the melanotroph versus corticotroph fate (Budry et al., 2012). Melanotrophs are quiescent (Langlais et al., 2013), in contrast with AL endocrine cells, however, they are particularly sensitive to mutations affecting cell cycle regulators (Quereda and Malumbres, 2009). In mice deleted for *p27*, encoding a protein best known for its negative cell cycle regulatory role (Sharma and Pledger, 2016), tumorigenesis in mature animals is the hallmark of pleiotropic effects (Fero et al., 1996, Kiyokawa et al., 1996),(Nakayama et al., 1996). Interestingly, this happens specifically in IL, while the anterior lobe is largely unaffected in mutants (Roussel-Gervais et al., 2010). In humans, *P27* mutations impairing protein function cause multiple endocrine neoplasia syndrome, MEN4, and low levels of P27 in tumours are generally associated with a poor prognosis. IL is residual in humans (Horvath et al., 1999). In patients affected by P27 mutations, AL pituitary tumors do occur, albeit rarely (Alrezk et al., 2017). In the developing murine pituitary, P27, along with P57, is necessary for cell cycle exit in differentiating endocrine cells (Bilodeau et al., 2009). However, its is not limited to cell cycle regulation, because in *Cyclin D1^-/-^;p27^-/-^* compound mutants, pituitary hyperplasia is still present (Tong and Pollard, 2001).

We have previously demonstrated that P27 can recruit co-repressors to regulate gene expression. More precisely, P27 can repress the expression of the transcription factor SOX2 during differentiation of induced pluripotent SC (iPSC) (Li et al., 2012). SOX2 is essential during embryonic development, for maintenance of adult SC properties in different cell lineages, and for somatic cell reprogramming (Sarkar and Hochedlinger, 2013). Its expression is elevated in many cancers and this is associated with increased proliferation, invasiveness, lineage plasticity, chemoresistance and, in consequence, poor prognosis (Wuebben and Rizzino, 2017),(Ku et al., 2017),(Mu et al., 2017). *In vivo*, the SOX2-P27 interaction is particularly relevant for IL because deletion of one copy of *Sox2* diminishes tumorigenesis in *p27^-/-^* animals (Li et al., 2012).

The cellular specificity of this interaction may be explained by the restricted pattern of expression of SOX2 in pituitary endocrine cells. Indeed, melanotrophs are the only endocrine cell type to express SOX2 (Goldsmith et al., 2016). However, SOX2 is also expressed in a population of adult pituitary SCs (Fauquier et al., 2008, Rizzoti et al., 2013, Andoniadou et al., 2013). Interestingly, in *p27^-/-^* mice, the thickness of the SOX2^+ve^ SC layer is increased (Li et al., 2012). Therefore, while the importance of the SOX2-P27 interaction for IL tumorigenesis is apparent, the modalities of this interaction and, in particular, the roles of SOX2 in the SCs and melanotrophs required for this are unknown.

In this study, we have examined how IL tumorigenesis develops in *p27^-/-^* mice, and explored the role of SOX2 in this process. As *p27^-/-^* IL become hyperplastic, melanotrophs tend to lose differentiated features and display increased levels of SOX2 expression, which is also observed in SCs. To determine the cellular origin of the tumors, and explore SOX2 function, we have performed lineage tracing experiments and deleted the gene separately in melanotrophs and SCs. Our results demonstrate that SOX2 is required in *p27^-/-^* melanotrophs, and also, independently, in SCs for tumorigenesis. Analyses of deletion of the *Sox2* regulatory region 2 (*Srr2*), which is the target of P27 repressive action, supports the pro-tumorigenic role of SOX2 in SCs. Finally, single cell transcriptomic analyses of IL reveals that activation of a SOX2-dependent MAPK pathway in SCs is important for tumorigenesis. In conclusion, our study emphasizes the pleiotropic effects of SOX2 in *p27^-/-^* IL and uncovers an unexpected role for SCs in IL tumour formation, suggesting that these, or the relevant signaling molecules, represent good anti-tumoral targets, in addition to tumour cells themselves.

## Results

### Characterization of intermediate lobe tumours in *p27^-/-^* pituitaries

We first examined P27 by immunofluorescence and observe widespread nuclear expression throughout the anterior pituitary with varying levels of expression (Fig 1A), in agreement with fluctuations of P27 according to the cell cycle (Sherr and Roberts, 1999). Anterior endocrine and stem cells proliferate rarely (Levy, 2002), (Fauquier et al., 2008) and P27 levels are relatively high in these. However, even though IL melanotrophs have been shown to be quiescent (Langlais et al., 2013), P27 immunostaining is much weaker compared with the AL and SCs. We then examined SOX2, which is expressed both in SCs (Fauquier et al., 2008), and, at lower levels, in melanotrophs (Goldsmith et al., 2016) (Fig 1A). In wild-type animals, P27 and SOX2 proteins are clearly co-expressed. Thus, despite the negative action of P27 on *Sox2* transcription both proteins can co-exist, probably reflecting the complex regulatory mechanisms controlling *Sox2* expression and/or availability of co-repressors recruited by P27. In *p27^-/-^* pituitaries, the SC layer appears thicker in mutants (Li et al., 2012). In agreement with a repressive action of P27 on *Sox2* expression (Li et al., 2012), SOX2/*Sox2* levels are significantly up-regulated both in SCs and melanotrophs in *p27^-/-^* pituitaries (Fig 1A-C).

**Figure 1.**
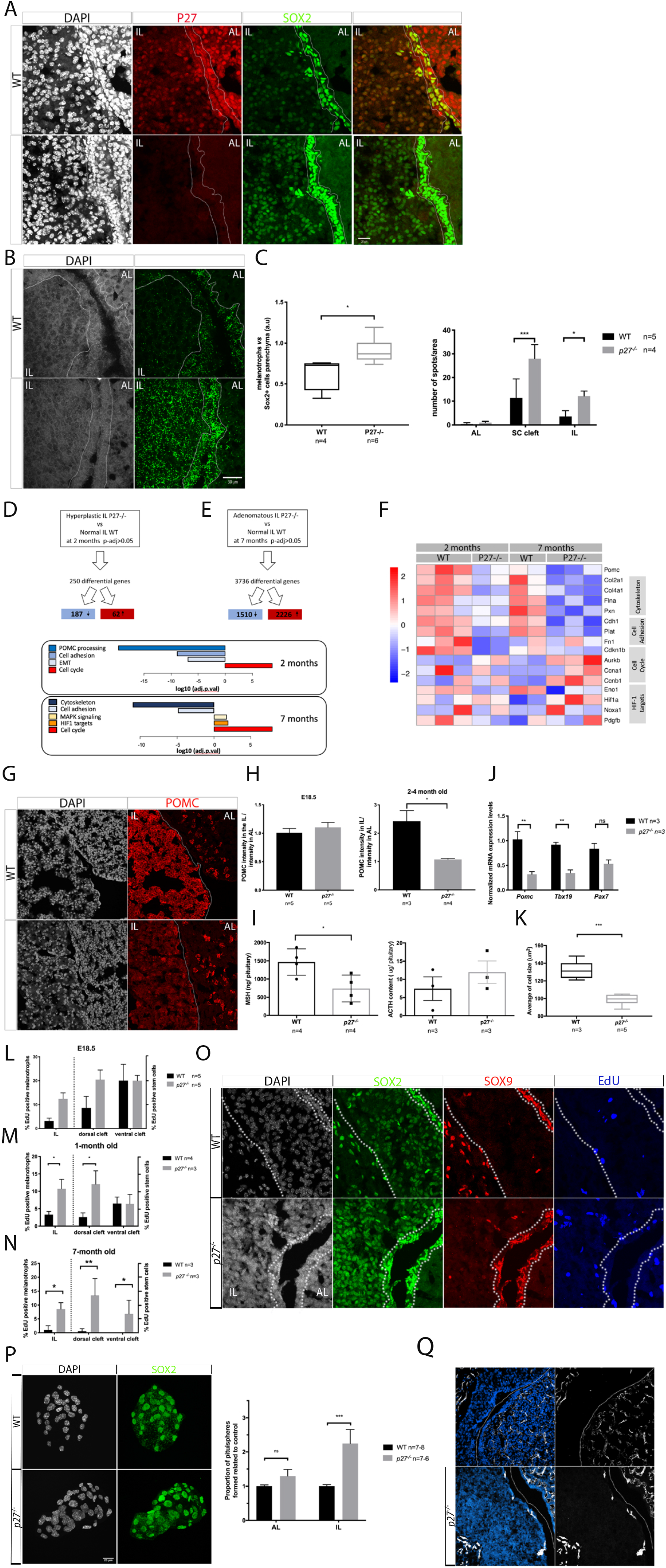
The expression of stem cell and differentiation markers is affected in *p27^-/-^* IL. **A)** Double immunofluorescence for p27 and SOX2 in wild-type and *p27^-/-^* adult pituitaries. p27 is expressed at high levels in AL and in SCs lining the cleft, but at low levels in melanotrophs. SOX2 is expressed at high levels in SCs and low levels in melanotrophs of control pituitaries. In *p27^-/-^* IL, SOX2 staining is more intense in both cell types. **B)** *In situ* hybridisation for *Sox2* in a 2-month old pituitary. In *p27^-/-^* IL higher levels of *Sox2* are observed in both melanotrophs and SCs. **C)** Quantification of *SOX2/Sox2* expression levels in *p27^-/-^* versus wild-type. Levels of expression were quantified after immunofluorescence in melanotrophs (A, left graph) and after *in situ* hybridisation, in the cell types indicated (B, right graph). In melanotrophs, the ratio between staining intensity in SOX2+melanotroph/SOX2+ SC in AL parenchyma is represented. In WT, SOX2 staining in melanotrophs is less intense than in AL SCs (0.63±0.20) while in *p27^-/-^* it becomes similar (0.91±0.16); (*p=0.019, n=4-6 in each group). Quantification after RNAscope show that in *p27^-/-^*, SCs flanking the cleft express *Sox2* at higher levels (28±6) than and wt (11.4±8, ***p=0.0002). The same is observed in melanotrophs: *p27^-/-^* (12±2) and wt (3.5±2.5, *p=0.0134), (n=5-4 in each group). **D-E)** Bulk RNAseq pathway analysis in 2 month-old IL before tumor formation (*p27^-/-^* n=2 vs wt n=3, adjusted p-value <0.05) (**D**), and in 7 month-old hyperplastic IL (*p27^-/-^* n=4 vs wt n=2) (**E**). **F)** Heatmap of representative genes expression from affected pathways shown in **D** and **E**. **G)** POMC immunofluorescence in control and *p27^-/-^* adult pituitaries. POMC staining appears reduced in mutant melanotrophs (IL), while corticotroph levels in AL appear normal. H) Graph representing quantification of POMC intensity in melanotrophs in relation to corticotrophs at e18.5 in wt (1.01±0.17, n=5) and *p27^-/-^* (1.10±0.19, n=5) and in 2 to 4 month-old animals, where levels are significantly reduced in mutants (wt:2.42±0.76, n=3; *p27^-/-^*:1.1±0.07, n=4, *p=0.0286). **I)** MSH and ACTH contents were measured by RIA in 2 month-old pituitaries. *p27^-/-^* pituitaries contain less MSH (*p=0.0306, n=4 in each group) while there is no significant difference in ACTH levels (right graph). **J)** Quantitative analysis of melanotroph marker expression levels by RT-qPCR in 2 to 3 month-old *p27^-/-^* and wt. *Pomc* (wt:1.03±0.27, n=3;*p27^-/-^*:0.32±0.09, n=3,**p=0.0013) and *Tbx19* levels (wt:0.94±0.08, n=3; *p27^-/-^*:0.30±0.10, n=3,**p=0.0071) were reduced in mutants. *Pax7* levels appeared similarly affected but this did not reach statistical significance (wt:0.84±0.19, n=3; *p27^-/-^*: 0.5±0.15, n=2). **K)** Average cell size of melanotrophs. *p27^-/-^* melanotrophs (99±5.6, n=5) are smaller compared to wt (132±10, n=3,***p=0.0007). **L-N)** Analysis of cell proliferation. EdU incorporation was quantified in melanotrophs, (EdU; SOX2;PAX7 triple positive/SOX2;PAX7 double positive in L or EdU; SOX2 double positive; SOX9 negative/SOX2 positive;SOX9 negative positive cells in M and N) and SCs (EdU;SOX2 double positive; PAX7 negative/SOX2 positive;PAX7 negative cells in L or EdU;SOX2;SOX9 triple positive /SOX2;SOX9 double positive in M and N). SCs were further distinguished as ventral or dorsal according to their localisation in the epithelial layer flanking the anterior or intermediate lobe respectively. At e18.5 (**L**) proliferation is not significantly different between mutants and controls (n=5), but there is a tendency toward more proliferation in mutant melanotrophs. In 1 month-old mice (**M**), there is a significant increase in cell proliferation in both *p27^-/-^* melanotrophs and dorsal cleft SCs (n = 3 to 4 mice/genotype, melanotrophs *p=0.0235, dorsal cleft *p=0.0419). In 7 month-old mice, (**N**) cell proliferation is increased in melanotrophs and both dorsal and ventral cleft SCs (n = 3 mice /genotype, melanotrophs *p=0.0167, dorsal cleft **p=0.0085, ventral cleft *p=0.0229). **O)** SOX2, SOX9 and EdU triple staining in 1-month old wt and *p27^-/-^* animals. **P)** Immunofluorescence for SOX2 on pituispheres obtained from IL wt and *p27^-/-^* 2 to 3 month-old animals. The proportion of pituispheres is increased in mutant IL (***p<0.0001, n=6-8 in each group). **Q)** Immunofluorescence for CD31 in 1-month old control and *p27^-/-^* pituitaries. In control pituitaries IL is avascular while *p27^-/-^* ILs show ectopic blood vessels at early stages (arrows). Scale bars represent 20μm for A and N and O, 30μm for B, 50μm for G, 100μm for P. Cleft is outlined in A, B, N and P, IL is outlined G, N.

To further characterize initiation and development of the tumours in *p27^-/-^* mice, we performed transcriptomic analyses from dissected IL (Fig 1D-F Supplementary Fig 1A). Before tumour formation, at two months of age, we detected 250 differentially expressed genes (DEG) between *p27^-/-^* and control samples. In seven-month old animals, when hyperplasia is very clear, there are 3736 DEG. At both stages, upregulation of cell cycle markers is observed, in agreement with the CDKI function of P27. We also observe downregulation of genes associated with the cytoskeleton and cell adhesion which may reflect disruption of the direct interaction between P27 and RhoA (Besson et al., 2004). Furthermore, downregulation of POMC and genes associated with its processing suggests an alteration of melanotroph function and/or identity in mutants.

We further investigated whether melanotroph function and/or identity have been altered, something that has not been reported previously (Fero et al., 1996, Kiyokawa et al., 1996),(Nakayama et al., 1996). We observe a significant downregulation of POMC/*Pomc* expression levels, exclusively in *p27^-/-^* melanotrophs post-natally, but prior to tumour formation (Fig 1G-H), while corticotrophs appear unaffected. In agreement with these data, radioimmunoassays (Borrelli et al.) show that levels of MSH are reduced while ACTH is unaffected in *p27^-/-^* pituitaries (Fig 1I). *p27^-/-^* mice are affected by gigantism, which has been attributed to general loss of cell cycle repression (Fero et al., 1996, Kiyokawa et al., 1996), (Nakayama et al., 1996), however, an additional explanation could be the significant increase in pituitary GH content we observe (Supplementary Fig 1B). This shows that the anterior pituitary is affected by loss of *p27*. TBX19 is a direct upstream regulator required for *Pomc* expression (Pulichino et al., 2003). In agreement with a reduction in *Pomc* levels, *Tbx19* levels are significantly reduced in IL, while *Pax7* (Budry et al., 2012) also appear reduced, although not to a statistically significant degree (Fig 1J). Moreover, *p27^-/-^* melanotrophs are smaller than those in wild-type pituitaries (Fig 1K). Altogether, these results suggest that melanotroph identity and/or post-natal maturation are altered in *p27^-/-^* mutants, and this occurs before tumours develop.

We then examined cell proliferation by quantifying incorporation of EdU in IL, dorsal (flanking IL), and ventral SC layer (flanking AL) (Fig 1L-O). In E18.5 embryos, proliferation tends to increase in both IL and its flanking SC layer in the mutants, however, this is not statistically significant (Fig 1L). In contrast, in one-month old mutant animals, we observe a significant increase in proliferation of both melanotrophs and adjacent SCs (Fig 1M-O). In seven-month old animals, increased proliferation is observed in all compartments of the mutants compared to controls (Fig 1N), in agreement with the transcriptomic analysis results (Fig 1E,F). To further test the proliferative capacity of SCs, we performed pituisphere assays (Fauquier et al., 2008) separately from anterior and intermediate lobe cells and observed an increased sphere forming ability from *p27^-/-^* IL cells (Fig 1P). Altogether these results suggest that, while both SC layers behave abnormally in *p27^-/-^* pituitaries, the SCs in direct contact with melanotrophs are more affected, suggesting that there may be some non-cell autonomous consequences to the loss of *p27*.

HIF-1 activation is associated with tumour angiogenesis and its targets are up-regulated in seven-month old *p27^-/-^* IL transcriptomes (Fig 1E). Moreover, *p27^-/-^* IL tumours are haemorrhagic (Fero et al., 1996). We thus examined CD31 expression and observe development of ectopic blood vessels in juvenile *p27^-/-^* IL, coincident with the abnormal increase in rates of cell proliferation (Fig 1Q).

In conclusion, *p27* deletion results in increased cell proliferation and ectopic vascularization in IL. While P27 starts to be expressed in differentiating embryonic endocrine cells (Bilodeau et al., 2009), defects consecutive to its loss appear to only become significant post-natally, both in melanotrophs and SCs. In addition, melanotroph identity and/or maturation appears affected in mutants.

### Characterization of the SOX2-P27 interaction in *p27^-/-^* mutants

Deletion of one copy of *Sox2* is sufficient to dramatically reduce occurrence of IL tumours (Fig 2A, C) (Li et al., 2012), however, hyperplasia is still observed in *p27^-/-^; Sox2^-/-^* IL (Fig 2A). We further characterized the SOX2-P27 interaction by demonstrating that survival of *p27^-/-^* mutants (12.5 months, n=28) was significantly improved by *Sox2* haploinsufficiency (17.1 months, n=10) (Fig 2B). However, in agreement with cell type specificity for the SOX2-P27 interaction, while pituitary and duodenum tumours were sensitive to *Sox2* dosage, lung tumour development was unaffected (Fig 2C).

**Figure 2.**
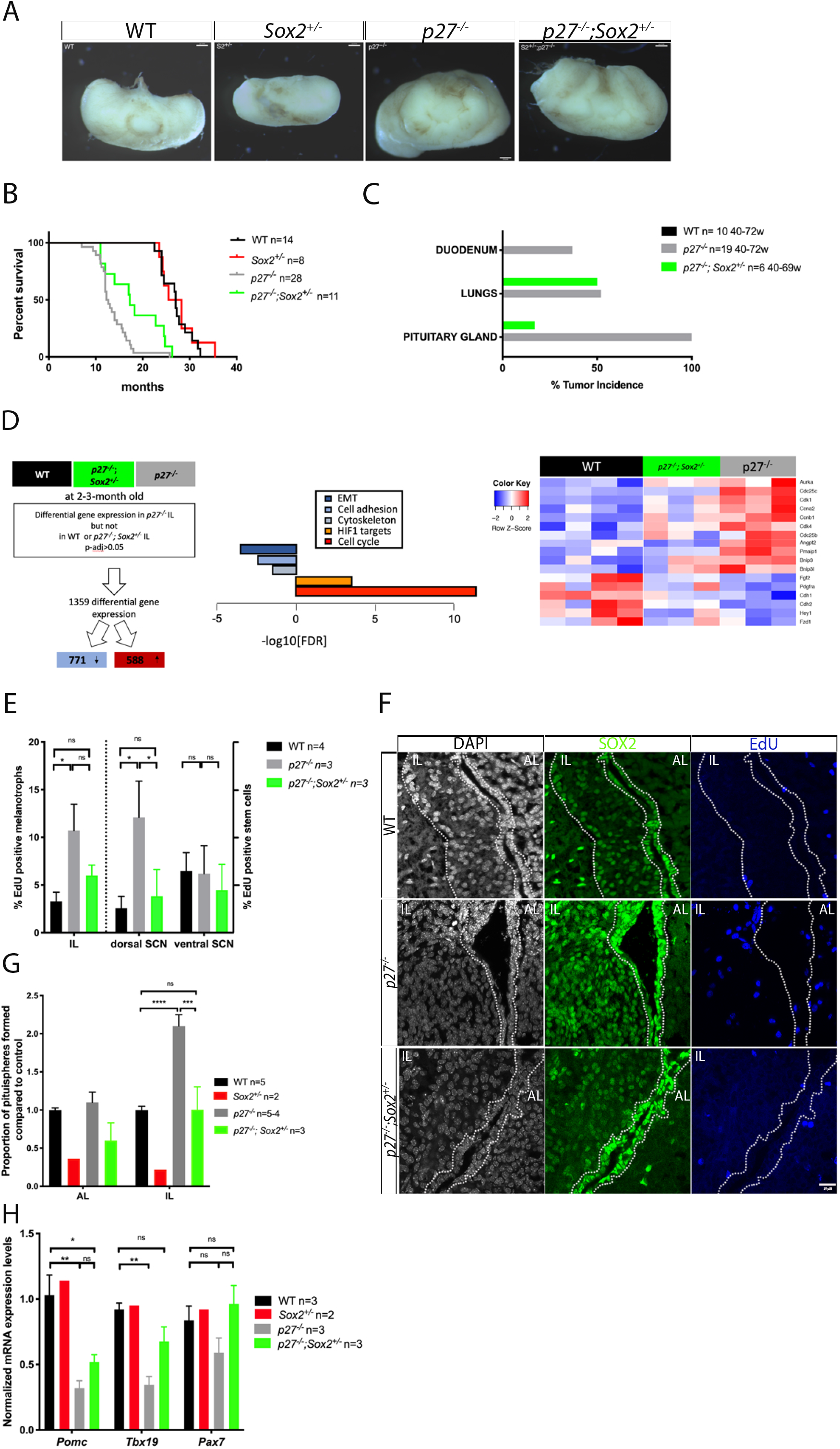
Deletion of one copy of *Sox2* in *p27^-/-^* animals improves survival and results in reduction of cell proliferation and impairment of tumorigenesis. **A)** Brightfield pictures of wt, *Sox2^+/-^, p27^-/-^ and p27^-/-^; Sox2^+/-^* pituitaries. In *p27^-/-^; Sox2^+/-^* IL tumourigenesis is impaired, but hyperplasia is observed. **B)** Kaplan-Meier survival curves for wt, *Sox2^+/-^; p27^-/-^* and *p27^-/-^; Sox2^+/-^* animals, *p27^-/-^; Sox2^+/-^* mutants survive longer (17.1 months, n=10) than *p27^-/-^* mutants (12.5 months, n=28, *p<0.01), but still significantly less than wt (27 months, n=14,****p<0.0001). **C)** Tumour incidence in the most affected organs in *p27^-/-^* mice. *Sox2* heterozygosity results in a reduction of tumour incidence in the pituitary and duodenum but not in lungs. **D)** Comparison modalities for RNA-seq analysis of wild-type, *p27^-/-^* and *p27^-/-^; Sox2^+/-^* 2 to 3 month-old IL samples (wt n=4, *p27^-/-^* n=3, *p27^-/-^; Sox2^+/-^* n=3, adjusted p-value <0.05) (left panel). Enriched pathways are represented (centre panel) and heatmap of representative genes expression from affected pathways (right panel). **E)** Analysis of cell proliferation. EdU incorporation was quantified in melanotrophs (EdU;SOX2 double positive; SOX9 negative/SOX2 positive;SOX9 negative cells) and SCs (EdU;SOX2;SOX9 triple positive/SOX2;SOX9 double positive cells). SCs were further distinguished as ventral or dorsal according to their localisation in the epithelial layer flanking the anterior or intermediate lobe respectively. Deletion of one copy of *Sox2* in compound mutants results in a reduction of cell proliferation compared to *p27^-/-^* samples in dorsal cleft SCs (n = 3-4 mice/genotype, *p=0.0208), melanotrophs. **F)** SOX2 and EdU double staining in 2-to 3-month old wt, *p27^-/-^* and *p27^-/-^; Sox2^+/-^* animals. There is a clear reduction in SOX2 levels in *p27^-/-^; Sox2^+/-^* IL. **G)** Proportion of pituispheres obtained from AL and IL wt, *Sox2^+/-^, p27^-/-^* and *p27^-/-^; Sox2^+/-^*, 2-to 3-month old animals. There is a reduction in the proportion of pituispheres formed from *p27^-/-^; Sox2^+/-^* IL compared to *p27^-/-^* samples, with the proportion in double mutants resembling that of wild type. wt (1±0.11,n=5), *Sox2^+/-^*(n=2), *p27^-/-^* (2.09±0.31,n=4) and *p27^-/-^;Sox2^+/-^* (1±0.52,n=3), ****p<0.0001 and ***p=0.0002; the reduction in *Sox2^+/-^* samples in agreement with the slightly hypomorphic pituitaries observed in *Sox2* heterozygous mice (73) and with SOX2 being required for proliferation of pituitary progenitors in the embryo (26) (74). **H)** Quantitative analysis of melanotroph marker expression levels by RT-qPCR in 2-to 3-month old wt, *Sox2*^+/-^, *p27^-/-^* and *p27^-/-^; Sox2*^+/-^ IL. In *p27^-/-^; Sox2^+/-^* animals, *Tbx19* expression levels are restored to wt levels. *Pomc*; wt (n=3) vs *p27^-/-^;Sox2^+/-^* (n=3), *p=0.03 and wt vs *p27^-/-^*(n=3), **p=0.0022, *Tbx19*; wt vs *p27^-/-^;Sox2*^+/-^ (n=3, p=ns) and wt vs *p27^-/-^* (n=3,*p=0.0169).

We then performed comparisons of the transcriptomes of wild-type, *p27^-/-^* and *p27^-/-^;Sox2^+/-^* IL (Fig 2D, Supplementary Fig 2A). To pinpoint genes underlying IL tumour impairment in compound mutants, and conversely those associated with higher levels of SOX2 and tumorigenesis, we focused our analyses on those that were exclusively differentially expressed in *p27^-/-^*, and therefore restored to control levels in *p27^-/-^;Sox2^+/-^* samples. We obtained a list of DEG and pathways comparable to our previous analysis (Fig 1D, Fig 2D). This suggests that SOX2 is not required for a specific aspect of tumour formation in *p27^-/-^* mice, but rather that it has a broad and therefore probably early role in tumour genesis. Reduced *Sox2* dosage is associated with a decrease in proliferation and HIF1-target gene expression, in agreement with less angiogenesis linked with reduced overproliferation in *p27^-/-^;Sox2^+/-^* mutants. We also observed an effect on genes involved in cytoskeleton, suggesting that reduction of *Sox2* allows restoration of a cellular phenotype resembling that of wild-type. Alternatively, alteration of cellular shape and contacts may be linked to variation in rates of cell proliferation.

We then analysed SOX2 expression, cell proliferation and pituisphere formation in *p27^-/-^;Sox2^+/-^* pituitaries (Fig 2E-G). There is a visible reduction in SOX2 levels in *p27^-/-^;Sox2^+/-^* IL (Fig 2F and Supplementary Fig 2B). Moreover EdU incorporation in compound mutants, both in melanotrophs and SCs is not significantly different from controls, in agreement with impaired tumour formation (Fig 2E). A similar effect is observed in pituisphere assays (Fig 2G). We moreover observe a reduction in pituisphere formation efficiency in *Sox2^+/-^* samples, suggesting a reduced proliferative capacity (Fig 2G). This is in agreement with the slightly hypomorphic pituitaries observed in *Sox2* heterozygous mice (Kelberman et al., 2006) and with SOX2 being required for proliferation of pituitary progenitors in the embryo (Goldsmith et al., 2016) (Jayakody et al., 2012).

Alteration of *p27^-/-^* melanotroph identity (Fig 1G) also appears to depend on SOX2 dosage, because levels of *Tbx19* which are reduced in *p27^-/-^* IL, are not significantly affected in *p27^-/-^;Sox2^+/-^* compared to control IL (Fig 2H and 5K). However, levels of *Pomc* were still significantly reduced in *p27^-/-^;Sox2^+/-^* IL compared to control.

### *Sox2 regulatory region 2 (Srr2)* mediates P27 effect on *Sox2* expression *in vivo*

Transcriptomic studies, proliferation and marker analyses all indicate that reduction of SOX2 dosage compromises most aspects of tumour genesis in *p27^-/-^* animals, implying an essential role for SOX2 in *p27^-/-^* melanotrophs and adjacent SCs for tumour development. To test whether the SOX2-P27 interaction relies solely on the *Srr2* enhancer (Li et al., 2012), we generated mice deleted for this region (Fig 3A, B). *Srr2^del/del^* mutants are viable (Supplementary Fig. 2C) and pituitary morphology is normal (Fig 3C). However, on a *p27^-/-^* background, the *Srr2* deletion prevents tumorigenesis, while hyperplasia of IL is still evident, as observed in *p27^-/-^;Sox2^+/-^* mutants (Fig 3C). In addition, the morphology of the retina, which is affected in absence of *p27* (Nakayama et al., 1996), is also improved in the *p27^-/-^; Srr2^del/del^* mutants. In particular, and again similarly to *p27^-/-^;Sox2^-/-^* mutants, the inner and outer layers have a reduced thickness and better organization compared to *p27^-/-^* retina (Fig 3D) (Li et al., 2012). In the pituitary, levels of SOX2 are reduced in both melanotrophs and SCs in *p27^-/-^; Srr2^del/del^* mutants compared to *p27^-/-^* samples (Fig 3F). Intriguingly, EdU incorporation analyses reveal a specific reduction of proliferation in SCs in compound mutants compared to *p27^-/-^* samples (Fig 3E). In agreement with this result, while cultures from *p27^-/-^* IL give rise to significantly more pituispheres than controls, the efficiency of pituisphere formation from *p27^-/-^; Srr2^del/del^* samples is reduced (Fig 3G). These analyses strongly suggest that *Srr2* has an exclusive role in mediating the SOX2-P27 interaction (see model Fig 3H). Moreover, the consequences of its deletion are more obvious in SCs, suggesting that *Srr2* plays a more important role in this compartment. Furthermore, this implies that SCs may be critical for IL tumorigenesis in *p27^-/-^* mutants.

**Figure 3.**
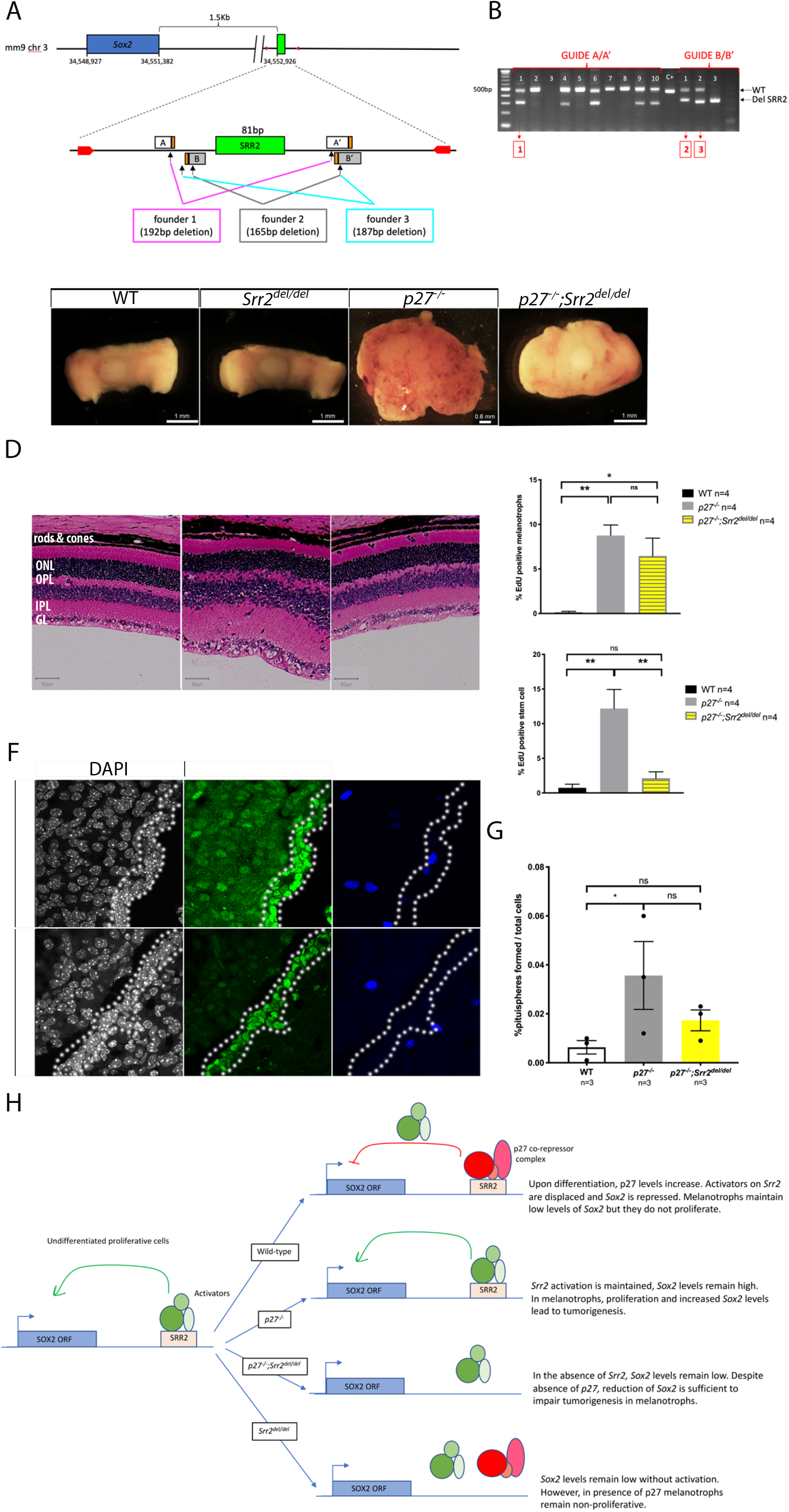
Deletion of *Srr2* in *p27^-/-^* animals results in reduction of cell proliferation in SCs and impairment of tumorigenesis. **A)** Schematic representation of the *Sox2* murine locus and *Srr2* deletions engineered by genome editing. Two pairs of sgRNA (pair 1: guide A_5’ and guide A’_3’ and pair 2: guide B_5’ and guide B’_3’, PAM sequence in orange) were designed for *Srr2* (green box) deletion. The primers used to genotype *Srr2* deleted animals are marked by red arrows. Three founders deleted for *Srr2* were used to generate stable lines (1 to 3); the size of the deletion is indicated for each line. **B)** Gel electrophoresis showing mosaic loss of *Srr2* in founder mice generated with sgRNA pair 1 (guide A/A’) or 2 (guide B/B’). The number above each lane corresponds to a individual founders. Boxed numbers below indicate strain derived from that particular founder. **C)** Brightfield pictures of wt, *Srr2^del/del^, p27^-/-^* and *p27^-/-^;Srr2^del/del^* pituitaries. Deletion of two copies of *Sox2* Srr2 on a *p27^-/-^* background is sufficient to delay tumorigenesis. **D)** Histological sections of H&E stained retinas. In *p27^-/-^;srr2^del/del^* retina, size and organization of the layers is improved compared to *p27^-/-^*. ONL = outer nuclear layer; OPL= outer plexiform layer, INL = inner nuclear layer, IPL=inner plexiform layer, GCL= ganglion cell layer. **E)** Analysis of cell proliferation. EdU incorporation was quantified in melanotrophs (EdU;SOX2 double positive;SOX9 negative/ SOX2 positive;SOX9 negative cells) and SCs (EdU;SOX2;SOX9 triple positive/SOX2;SOX9 double positive cells) in 7-month old wt, *p27^-/-^* and *p27^-/-^;Srr2^del/del^* animals. In *p27^-/-^;Srr2^del/del^* animals, there is a reduction of proliferation in stem cells compared to *p27^-/-^*(*p27^-/-^* (12.33±5.37, n=4) vs *p27^-/-^;Srr2^del/del^* (2.11±1.88,n=4), **p=0.0061), but not in melanotrophs (*p27^-/-^* (8.75±2.36) vs *p27^-/-^;Srr2^del/del^* (6.45±4.01), p=ns). **F)** SOX2 and EdU double staining in 7-month old *p27^-/-^* and *p27^-/-^;Srr2^del/del^* animals. SOX2 expression appears decreased *in p27^-/-^;Srr2^del/del^* IL. Scale bars represent 100mm for A, 20μm for F and O, 1-0.6mm for L, 50μm for M. Cleft is underlined in F and O, IL is underlined in F wt panel. **G)** Proportion of pituispheres obtained from IL in wt, *p27^-/-^* and *p27^-/-^; Srr2^del/del^*. The percentage of spheres formed from *p27^-/-^; Srr2^del/del^* (0.02±0.007, n=3, p=ns) is not significantly higher than wt (0.006±0.005, n=3), in contrast with *p27^-/-^* samples (0.04±0.02, n=3, *p=0.0344) demonstrating that loss of *Srr2* has an impact on sphere forming efficiency in mutants. **H)** Model illustrating the effects of p27 on Srr2 enhancer and consequences of their loss in melanotrophs. Srr2^del/del^ animals do not phenocopy aspects of the p27^-/-^ phenotype, which might have been expected if all the enhancer does is mediate repression of Sox2 by P27. However, in agreement with the phenotype of p27^-/-^; Srr2^del/del^ animals, deletion of Srr2 prevents both repression and derepression of SOX2.

### Requirement for SOX2 in *p27^-/-^* melanotrophs

Conditional deletion of *p27* in melanotrophs results in formation of tumours, therefore transformation is cell-autonomous in this context (Chien et al., 2006). To better characterise the role of SOX2 for tumour genesis, we deleted one copy of the gene exclusively in melanotrophs in *p27^-/-^* mutants. We initially used *Pomc-CreERT2* which displays a mosaic pattern of recombination in IL (Fig 4A). To look at heterozygosity for *Sox2*, we induced Cre activity in four-week old *Pomc-CreERT2;Sox2^fl/+^;p27^-/-^* animals and controls, which were then examined six to eight months later. Tumour development was unaffected by the mosaic deletion of one copy of *Sox2* in *p27^-/-^* melanotrophs (Fig 4B). In agreement with this observation, proliferation was unaffected by *Sox2* reduction, as we observed a similar increase in EdU incorporation in *Pomc-CreERT2;Sox2^fl/+^;p27^-/-^* and *Pomc-CreERT2;p27^-/-^* IL compared to controls (Fig 4B,C). To analyse more directly the effect of the loss of one copy of *Sox2* on melanotrophs proliferative ability, we analysed EdU incorporation in *Pomc-CreERT2;Sox2^fl/+^;p27^-/-^;Rosa26^ReYEP^* samples. We compared the percentages of eYFP;PAX7; EdU triple positive and eYFP negative, PAX7; EdU double positive melanotrophs, assuming that *Rosa26^ReYEP^* recombination also reflected *Sox2* heterozygous deletion. We observed a significant reduction of EdU incorporation in eYFP positive versus negative cells (Fig 4D). Therefore, reduction of SOX2 dosage in *p27^-/-^* melanotrophs diminishes their proliferative capacity.

**Figure 4.**
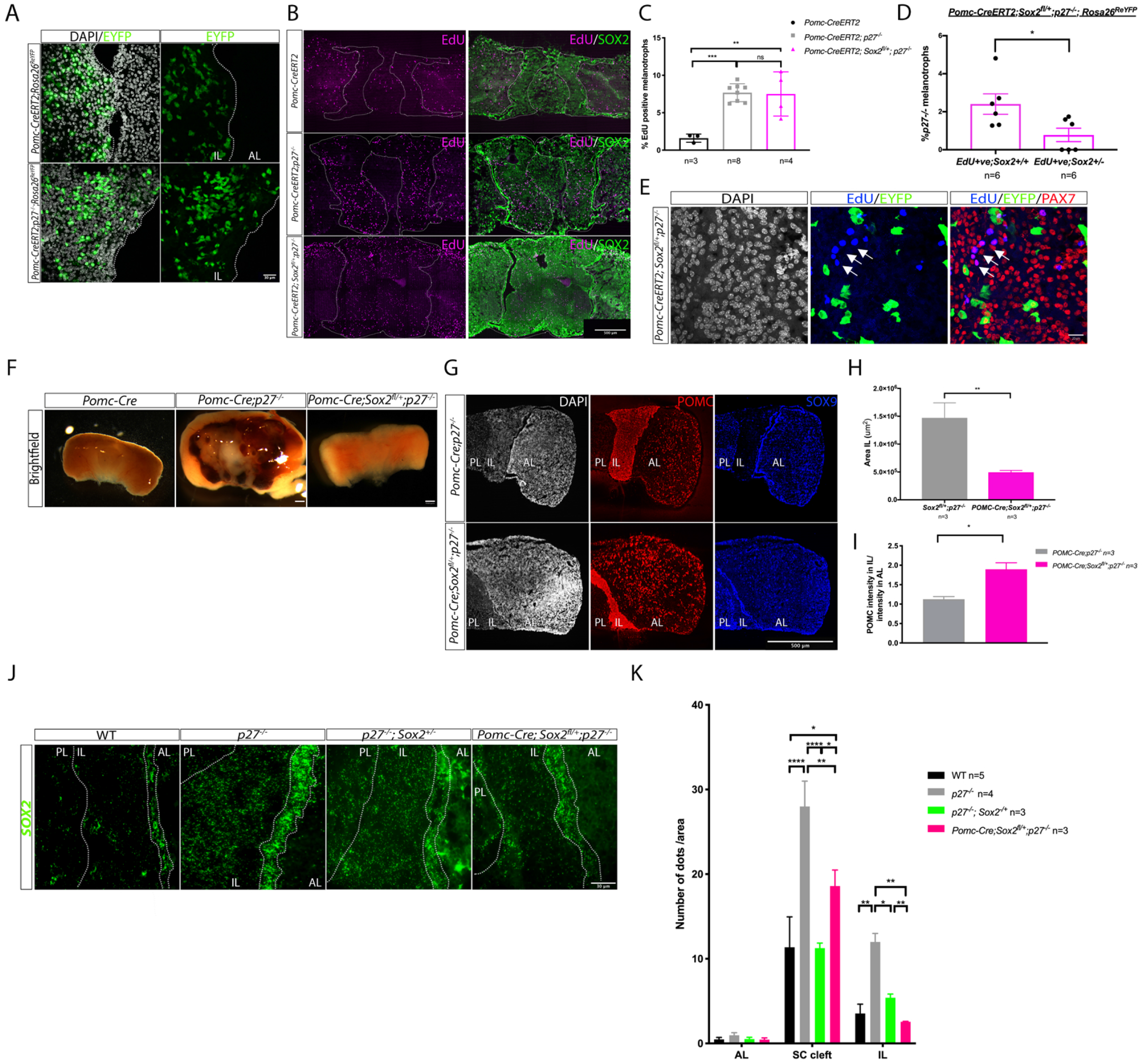
Deletion of one copy of *Sox2* in *p27^-/-^* melanotrophs prevents IL tumorigenesis. **A)** eYFP mosaic recombination pattern in 6-month old *Pomc-CreERT2;Rosa26R^eYEP/+^* and *Pomc-CreERT2;p27^-/-^;Rosa26R^eYEP/+^* IL. Only a proportion of melanotrophs are eYFP positive after tamoxifen treatment. **B)** SOX2 and EdU double staining in 6-to 8-month old *Pomc-CreERT2, Pomc-CreERT2;p27^-/-^ and Pomc-CreERT2:Sox2^fl/+^;p27^-/-^* pituitaries. **C)** Analysis of cell proliferation. EdU incorporation was quantified in melanotrophs (EdU;SOX2 low;POMC triple positive/DAPI nuclei in IL). EdU levels are similarly elevated in both *p27^-/-^* samples. *Pomc-CreERT2:Sox2^fl/+^;p27^-/-^* (7.53±2.92,n=4, **p=0.0023) and *Pomc-CreERT2;p27^-/-^*(7.68±1.20,n=8, ***p=0.0007) vs *Pomc-CreERT2* (controls), (1.60±0.55, n=3). **D-E)** EdU incorporation was quantified in *Sox2^+/+^* melanotrophs (identified as PAX7+ve;eYFP-ve) (2.40±1.31, n=6) and in *Sox2*^+/-^ melanotrophs (identified as PAX7;eYFP double positive) (0.78±0.86, n=6) in 2.5-to 6-month old *Pomc-CreERE2:Sox2^fl/+^;p27^-/-^:Rosa26^ReYEP^* mice, assuming that *Rosa26^ReYEP^* recombination reflects *Sox2* heterozygous deletion. In *Sox2*^+/-^ melanotrophs EdU incorporation is significantly reduced compared to *Sox2^+/+^* ones (*p=0.0210) demonstrating that loss of one copy of *Sox2* results in reduced proliferation in *p27^-/-^* melanotrophs. **E)** EdU, eYFP and PAX7 triple staining. Most EdU;PAX7 positive cells (arrows) do not express eYFP in *Pomc-CreERT2;Sox2^fl/+^;p27^-/-^;Rosa26^ReYEP^* IL. **F)** Brightfield pictures of *Pomc-Cre, Pomc-Cre;p27^-/-^* and *Pomc-Cre;Sox2^fl/+^;p27^-/-^* pituitaries. Deletion of one copy of *Sox2* using Pomc-Cre prevents IL tumorigenesis. **G)** Double immunofluorescence for POMC and SOX9 in 6-to 10-month old animals illustrating reduced IL size in *Pomc-Cre:Sox2^fl/+^;p27^-/-^*. **H)** Measurement of IL area in *Pomc-Cre;p27^-/-^* (1472054±808382, n=3) and Pomc-Cre;Sox2^fl/+^;p27^-/-^ (495612±185861, n=3). IL area *in Pomc-Cre;Sox2^fl/+^;p27^-/-^* pituitaries represents a third of IL area in *p27^-/-^* mutants (**p=0.0041). **I)** Quantification of POMC intensity in melanotrophs in relation to corticotrophs in 6-to 10-month old *Pomc-Cre;p27^-/-^* (1.13±0.11,n=3) vs *Pomc-Cre;Sox2^fl/+^;p27^-/-^* (1.89±0.29,n=3) animals, where levels are significantly increased, *p=0.0127. **J)** *In situ* hybridisation for *Sox2* in 6.25-to 10-month old wild-type, *p27^-/-^*, *p27^-/-^; Sox2*^+/-^ and *Pomc-Cre;Sox2^fl/+^;p27^-/-^* animals. **K)** Quantification of *Sox2* levels following *in situ* hybridisation in AL, SC and IL. *Sox2* expression levels are reduced in both *p27^-/-^; Sox2*^+/-^ and *Pomc-Cre;Sox2^fl/+^;p27^-/-^* melanotrophs and stem cells compared to *p27^-/-^*, showing respectively cell autonomous and tumor-inducing effects following *Sox2* deletion in melanotrophs. Quantification of *Sox2* levels in melanotrophs: *p27^-/-^* (12.17±2.15,n=4), wt (3.5±2.5, **p<0.003), *p27^-/-^;Sox2^+/-^* (5.4±1, *p=0.0276), *Pomc-Cre;Sox2^fl/+^;p27^-/-^* (2.56±0.1, **p=0.0038). Quantification of *Sox2* levels in SCs: *p27^-/-^* (28±6,n=4), wt (11.4±8,n=5, ****p<0.0001), *p27^-/-^;Sox2^+/-^* (11.3±1.4,n=3, ****p<0.0001), *Pomc-Cre;Sox2^fl/+^;p27^-/-^* (19±3,n=3, **p=0.0039). Scale bars represent 30 μm for A and J, 20 μm for E, 500 μm for B, F and G. IL is underlined A and B. IL and stem cell cleft underlined in I.IL=intermediate lobe and AL=anterior lobe.

We then hypothesized that a more efficient Cre-driver may block tumour formation; we therefore generated *Pomc-Cre;Sox2^fl/+^;p27^-/-^* animals. We showed previously that SOX2 is required for acquisition of melanotroph identity, however homozygous deletion of the gene once melanotrophs have up-regulated PAX7 does not have an apparent effect on cell fate acquisition (Goldsmith et al., 2016). There were no IL tumours or signs of hyperplasia in any four-to seven-month old *Pomc-Cre;Sox2^fl/+^;p27^-/-^* animals (Fig 4F-H). Moreover POMC staining intensity was increased in *Pomc-Cre;Sox2^fl/+^;p27^-/-^* melanotrophs compared to *p27^-/-^* samples (Fig 4I), indicating restoration of a normal melanotroph phenotype. This demonstrates that SOX2 is required in melanotrophs for *p27* deletion to result in tumourigenesis. Strikingly, the morphology of IL appears completely normal in *Pomc-Cre;Sox2^fl/+^;p27^-/-^* (Fig 4F,G) compared to *p27^-/-^;Sox2^+/-^* IL where hyperplasia is clear (Fig 2A). This difference in phenotype is unexpected, because in both mutants the genotype of melanotrophs is the same: only one allele of *Sox2* is active while *p27* is deleted. However, different regulatory mechanisms may affect the levels of expression of the remaining copy of *Sox2* in *Pomc-Cre;Sox2^fl/+^;p27^-/-^* versus *p27^-/-^;Sox2^+/-^* melanotrophs. We thus quantified *Sox2* levels and observe an up-regulation in *p27^-/-^*, and to a lesser extent in *p27^-/-^;Sox2^+/-^* samples (Fig 4J,K). In agreement with the absence of hyperplasia, we observed a further reduction in *Sox2* expression in *Pomc-Cre;Sox2^fl/+^;p27^-/-^* versus *p27^-/-^;Sox2^+/-^* samples, suggesting that regulation of *Sox2* is indeed modulated according to the spatiotemporal pattern of its deletion. Furthermore we also observe, albeit to a lesser extent, reduction of *Sox2* expression in *Pomc-Cre;Sox2^fl/+^;p27^-/-^* SCs.

In conclusion, while cell transformation in *p27^-/-^* melanotrophs is known to be cell-autonomous (Chien et al., 2006), we demonstrate here that SOX2 is required in these cells for tumour development in *p27^-/-^* mutants. Moreover, reduction of *Sox2* expression in SCs in *Pomc-Cre:Sox2^fl/+^;p27^-/-^* suggests a non cell autonomous role, and implies an interaction between melanotrophs and SCs.

### SOX2 is required independently in SCs for induction of melanotroph tumours in *p27^-/-^* animals

Because deletion of *p27* induces up-regulation of SOX2 in SCs, we ask whether the SCs flanking the IL directly give rise to tumour cells or induce tumour formation. Given that, in the context of the pituitary, SOX9 is expressed uniquely in the SCs (27), we first performed lineage tracing experiments using *Sox9^iresCreERT2^* (Furuyama et al., 2010). Cre activity was induced in four-week-old *Sox9^iresCreERT2/+^;Rosa26^ReYEP/+^* animals and pituitaries harvested six to eight months later. In controls, the vast majority of eYFP positive cells are SOX9 positive SCs (Rizzoti et al., 2013) (Fig 5A). In agreement with a low turnover of melanotrophs (Langlais et al., 2013), very few eYFP positive cells are present in IL (Fig 5A); these are negative for SOX9 and PAX7, where the latter indicates they are not melanotrophs. Their identity is unclear; they may originate from the pituitary SC layer or the posterior lobe where SOX9 is also expressed. In *p27^-/-^* animals we also observe that most eYFP positive cells are SOX9 positive SCs. However, in contrast with *Sox9^iresCreERT2/+^;Rosa26^ReYEP/+^* controls, we observe some rare eYFP;PAX7 positive melanotrophs. These results clearly demonstrate that SCs are not at the origin of the melanotroph tumours in *p27^-/-^* animals. However, tumour formation and/or lack of *p27* in SCs, drives commitment of rare SCs toward the melanotroph lineage.

**Figure 5.**
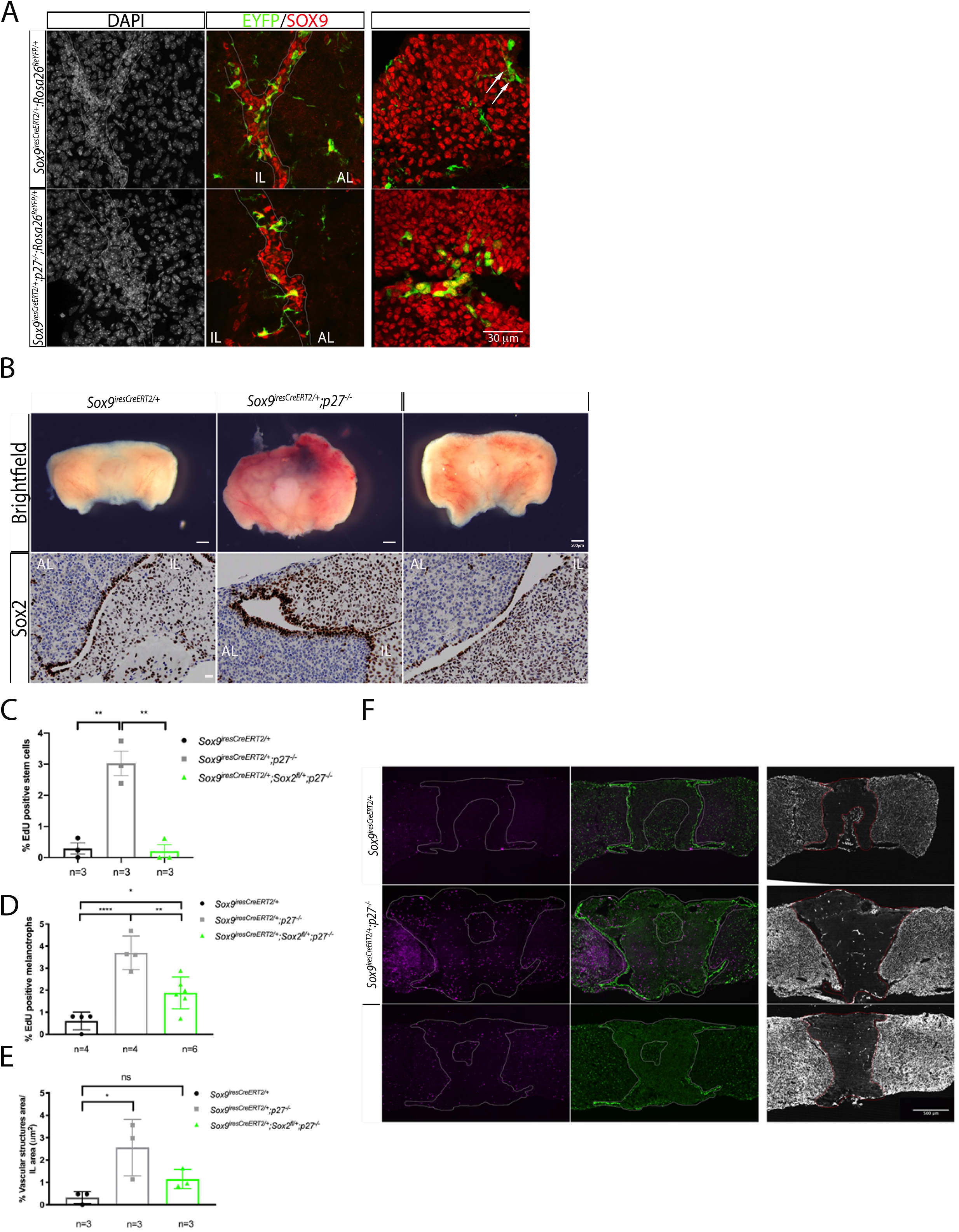
SOX2 is required in SCs for tumorigenesis in *p27^-/-^* IL. **A)** eYFP, SOX9 (middle panel) and eYFP, PAX7 (right panel) double stainings in 7-to 12-month old *Sox9^iresCreERT2/+^;Rosa26^ReYEP/+^* and *Sox9^iresCreERT2/+^;p27^-/-^;Rosa26^ReYEP/+^* animals. eYFP;SOX9 double positive cells are localized in the epithelium lining the cleft in both genotypes (arrows). Very few eYFP+, SOX9- and PAX7-cells are present in *Sox9^iresCreERT2/+^;Rosa26^ReYEP/+^* animals (arrows). In *Sox9^iresCreERT2/+^;p27^-/-^;Rosa26^ReYEP/+^* pituitaries some rare eYFP;PAX7 double-positive cells are present, demonstrating low levels of differentiation into melanotroph (left panel). **B)** Brightfield pictures (upper panel) and SOX2 immunohistochemistry (bottom panel) on sections of *Sox9^iresCreERT2/+^;Rosa26^ReYEP/+^, Sox9^iresCreERT2/+^;p27^-/-^* and *Sox9^iresCreERT2/+^;Sox2^fl/+^;p27^-/-^* pituitaries. In *Sox9^iresCreERT2/+^;Sox2^fl/+^;p27^-/-^* pituitaries tumorigenesis is impaired (one-year old 14% tumour incidence, n=7) compared to *Sox9^iresCreERT2/+^;p27^-/-^* (one-year old, n=5, 100% tumour incidence). The thickness of the SOX2 positive SC epithelium resembles that of control, demonstrating that reduction of *Sox2* dosage in SCs impairs *p27^-/-^* ILtumorigenesis. **C,D,F)** Analysis of cell proliferation. EdU incorporation was quantified in SCs and melanotrophs in 7-to 12-month old *Sox9^iresCreERT2/+^, Sox9^iresCreERT2/+^;p27^-/-^* and *Sox9^iresCreERT2/+^, Sox9^fl/+^;p27^-/-^* IL. **C)** In SCs (EdU;SOX2;SOX9 triple positive/SOX2;SOX9 double-positive cells), there is a significant reduction of proliferation in *Sox9^iresCreERT2/+^, Sox9^fl/+^;p27^-/-^* samples (0.20±0.35, n=3, **p=0.0011) compared to *Sox9^iresCreERT2/+^;p27^-/-^* ones (3±0.68, n=3). In fact, proliferation in *Sox9^iresCreERT2/+^, Sox9^fl/+^;p27^-/-^* SC is similar to *Sox9^iresCreERT2/+^* (0.28±0.32,n=3, p=ns). **D)** In melanotrophs (POMC positive cells in IL/DAPI positive nuclei), there is a reduction of cell proliferation in *Sox9^iresCreERT2/+^, Sox9^fl/+^;p27^-/-^* samples (1.88±0.72, n=6, **p=0.0033) compared to *Sox9^iresCreERT2/+^;p27^-/-^* (3.7±0.76, n=4); again proliferation in *Sox9^iresCreERT2/+^, Sox9^fl/+^;p27^-/-^* is similar to *Sox9^iresCreERT2/+^* controls (0.60±0.4,n=4, *p=0.03) **E)** Quantification of the vascular structures ectopic development in 7-to 12-month old. *Sox9^iresCreERT2/+^, Sox9^iresCreERT2/+^;p27^-/-^* and *Sox9^iresCreERT2/+^, Sox9^fl/+^;p27^-/-^* animals. Deletion of one copy of *Sox2* in SC leads to a reduction in the development of ectopic blood vessels in IL. IL vascular structures area: *Sox9^iresCreERT2/+^* (0.32±0.28, n=3), *Sox9^iresCreERT2/+^;p27^-/-^* (2.56±1.26, n=3, (*p<0.03)) and *Sox9^iresCreERT2/+^, Sox9^fl/+^;p27^-/-^* (1.15±0.43, n=3). **F)** EdU, SOX2 double staining in 7-to 12-month old *Sox9^iresCreERT2/+^, Sox9^iresCreERT2/+^;p27^-/-^* and *Sox9^iresCreERT2/+^, Sox9^fl/+^;p27^-/-^* pituitary glands. Right panel, immunofluorescence for CD31 showing ectopic blood vessel formation reduction in *Sox9^iresCreERT2/+^, Sox9^fl/+^;p27^-/-^* sample. Scale bars represent 30 μm for A, 20 μm for B (bottom panel), 500 μm for B (upper panel), F. IL is underlined F. Stem cell cleft underlined in A.IL=intermediate lobe and AL=anterior lobe.

To investigate the role of SOX2 in SCs during tumourigenesis one allele of the gene was removed exclusively in SCs in *Sox9^iresCreERT2/+^;Sox2^fl/+^;p27^-/-^* animals. Cre activity was induced in four-week old animals and pituitaries harvested when the animals were six-to twelve-month old. Strinkingly, only one amongst seven 12-month old *Sox9^iresCreERT2/+^;Sox2^fl/+^;p27^-/-^* animals developed an IL tumour, while all *p27^-/-^* controls were affected (Fig 5B, upper panel). The IL is however consistently hyperplastic in *Sox9^iresCreERT2/+^;Sox2^fl/+^;p27^-/-^* animals, similarly to *p27^-/-^;Sox2^+/-^* mice. These results unequivocaly demonstrate that SOX2 is required independently in SCs for tumours to develop in *p27^-/-^* IL. The thickness of the SOX2 positive SC layer appeared reduced in *Sox9^iresCreERT2/+^;Sox2^fl/+^;p27^-/-^* compared to *Sox9^iresCreERT2/+^;p27^-/-^* (Fig 5B, lower panel), and there were fewer SCs upon *Sox2* reduction (Supplementary Fig 3A). SOX2 staining intensity in SC was also reduced (Fig 5F, Supplementary Fig 3B). Furthermore, we observed a significant reduction of proliferation in both SC and melanotrophs in six-month old *Sox9^iresCreERT2/+^;Sox2^fl/+^;p27^-/-^* IL compared to *Sox9^iresCreERT2/+^;p27^-/-^* samples (Fig 5C, D, F). In addition, formation of ectopic vascular structures is significantly decreased (Fig 5E, F).

Altogether these results indicate that, while SCs are not themselves giving rise to melanotroph tumours in *p27^-/-^* IL, SOX2 activity in the SCs, and consequently the SCs themselves, are required for tumourigenesis in *p27^-/-^* IL.

### Characterization of the transcriptional consequences of altering *Sox2* levels in *p27^-/-^* IL SCs

To analyse the role of SOX2 in SC, we performed single cell RNA sequencing analysis (scRNAseq) on ILs from three-month old *Sox9^iresCreERT2/+^, Sox9^iresCreERT2/+^;p27^-/-^* and *Sox9^iresCreERT2/+^;Sox2^fl/+^;p27^-/-^* animals, where Cre activity had been induced at birth. Dataset clustering was performed by generating UMAP plots (Fig 6A-B). The three datasets were integrated and clusters identified according to the expression of known DEG, and by comparison with published studies (Cheung et al., 2018b, Mayran et al., 2019). The major cell cluster corresponds to melanotrophs as expected (*Tbx19, Pomc* and *Pax7* positive). The SC cluster (*Sox2*, *Sox9* positive) is proportionally larger in our IL dataset in comparison with studies performed on the whole pituitary (Cheung et al., 2018b, Mayran et al., 2019). We independently re-analysed the SC (Fig 6C, D) and melanotroph clusters (Fig 6G, H). Within each cell type we observed that cells were mostly, but not perfectly, clustered according to their genotype. We thus aimed at examining DEGs between the clusters to characterise the direct effect of *Sox2* dosage on SCs, and the consequences on melanotrophs. More precisely, we analysed DEGs in *Sox9^iresCreERT2/+^;p27^-/-^* compared to *Sox9^iresCreERT2/+^* and *Sox9^iresCreERT2/+^;Sox2^fl/+^;p27^-/-^*.

**Figure 6.**
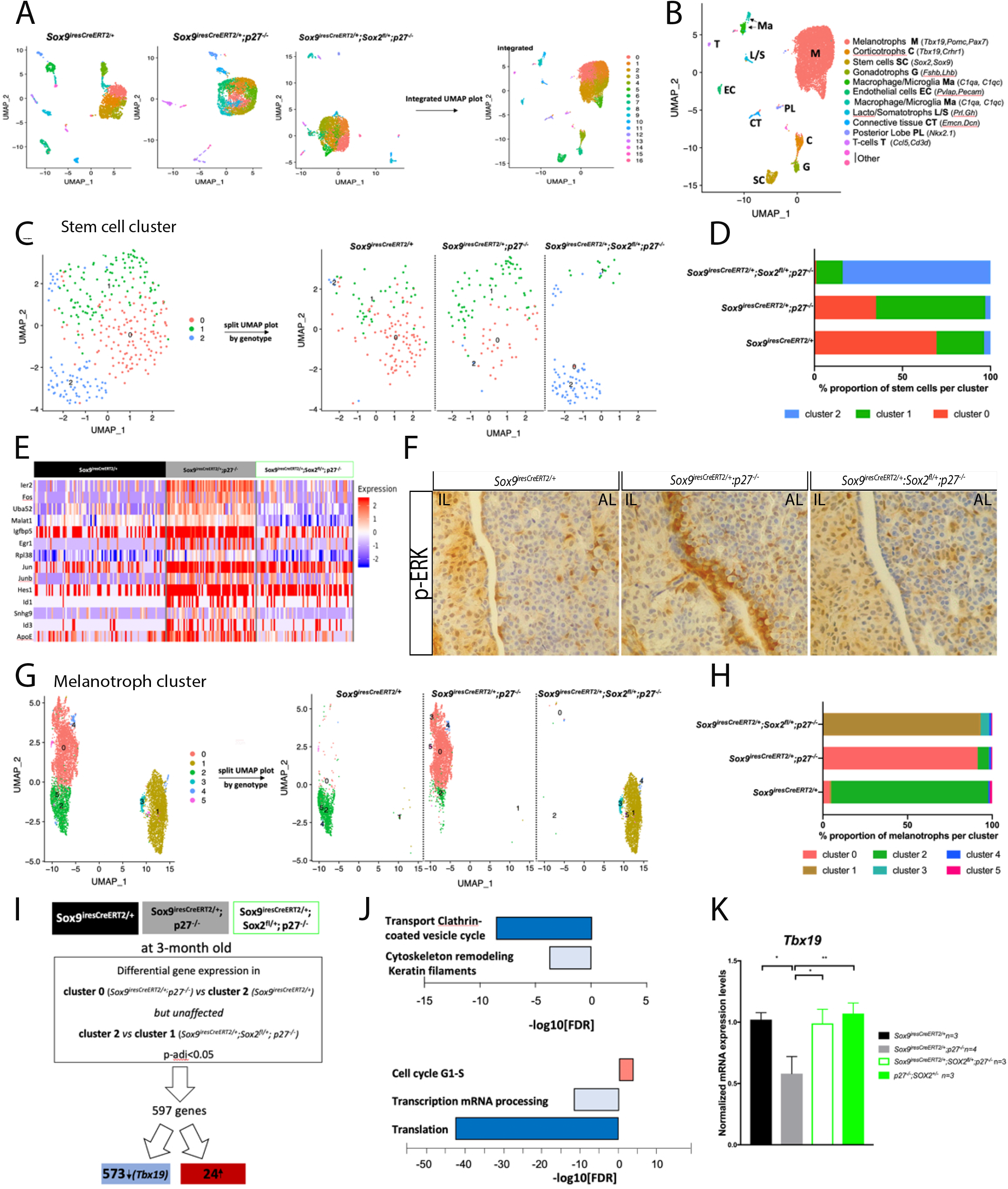
Characterization of the transcriptomic effects of *Sox2* dosage reduction in SCs in *p27^-/-^* IL by single cell analysis. **A)** UMAP (Uniform Manifold Approximation and Projection) visualization of 3 month-old *Sox9^iresCreERT2/+^* (n=2110 cells), *Sox9^iresCreERT2/+^:p27^-/-^* (n=4527cells) and *Sox9^iresCreERT2/+^;Sox2^fl/+^;p27^-/-^* (n=4305 cells) single cell RNAseq datasets from dissected ILs. Individual datasets were integrated on a unique UMAP plot from the three different genotypes (right panel). **B)** Identification of clusters according to the expression of known markers. Melanotroph clusters, identified by the expression of *Pomc, Tbx19* and *Pax7* genes and non segregated in PCA, were merged in one single cluster. **C)** Subclustering of the stem cell fraction (0.9 resolution) shows that segregation only partially correlates with genotype (right panel). **D)** Representation of the proportion of cells of the indicated genotype in each cluster, *Sox9^iresCreERT2/+^* (n= 137 cells), *Sox9^iresCreERT2/+^:p27^-/-^* (n= 104 cells) and *Sox9^iresCreERT2/+^;Sox2^fl/+^;p27^-/-^* (n= 82 cells). **E)** Heatmap showing expression of 14 genes selected out of 36 differentially expressed genes (adjusted p-value <0.05) in *Sox9^iresCreERT2/+^:p27^-/-^* compared with *Sox9^iresCreERT2/+^* or *Sox9^iresCreERT2/+^;Sox2^fl/+^;p27^-/-^*. **F)** Immunohistochemistry for p-ERK on *Sox9^iresCreERT2/+^*, *Sox9^iresCreERT2/+^:p27^-/-^* and *Sox9^iresCreERT2/+^;Sox2^fl/+^;p27^-/-^* pituitaries. While the MAPK/ERK pathway seems overactive in *p27^-/-^* SCs, p-ERK levels returns to normal levels upon deletion of one copy of *Sox2* in SCs. **G)** Subclustering of the melanotroph fraction (0.2 resolution) shows that segregation correlates with genotype (right panel). **H)** Representation of the proportion of cells of the indicated genotype in the most abundant cluster, *Sox9^iresCreERT2/+^*(n= 1272 cells), *Sox9^iresCreERT2/+^:p27^-/-^* (n= 3561 cells) and *Sox9^iresCreERT2/+^;Sox2^fl/+^;p27^-/-^* (n= 3006 cells). **I)** Strategy followed for scRNA-seq analysis of *Sox9^iresCreERT2/+^:p27^-/-^* vs *Sox9^iresCreERT2/+^* or/and *Sox9^iresCreERT2/+^;Sox2^fl/+^;p27^-/-^* melanotrophs. **J)** Upper panel, pathway analysis on genes differentially expressed (adjusted p-value <0.05) in melanotrophs following the strategy delineated in (I). Pathways associated with secretory function and cytoskeleton are downregulated in *Sox9^iresCreERT2/+^:p27^-/-^* cells. Bottom panel, process network analysis performed on genes differentially expressed between *Sox9^iresCreERT2/+^:p27^-/-^* and *Sox9^iresCreERT2/+^;Sox2^fl/+^;p27^-/-^* melanotrophs (adjusted p-value <0.05). Processes associated with translation and transcription are downregulated in *Sox9^iresCreERT2/+^:p27^-/-^* melanotrophs, while genes promoting cell cycle progression are upregulated. **K)** RT-qPCR analysis of *Tbx19* levels in 5 to 12 month-old *Sox9^iresCreERT2/+^*, *Sox9^iresCreERT2/+^:p27^-/-^*, *Sox9^iresCreERT2/+^;Sox2^fl/+^;p27^-/-^* and *p27^-/-^;Sox2*^+/-^ IL. *Tbx19* levels are increased in *Sox9^iresCreERT2/+^;Sox2^fl/+^;p27^-/-^* (0.99±0.20, n=3, *p=0.02) and *p27^-/-^*:*Sox2*^+/-^ (1.07±0.15, n=3, **p=0.0044) vs *Sox9^iresCreERT2/+^:p27^-/-^* (0.58±0.28, n=4).

In SCs, (Fig 6C, D) thirty-six DEGs were identified by comparing subcluster 1 (which has the highest proportion of *p27^-/-^* cells) to the two other subclusters (Supplementary Table 1). Thirty of the DEGs were upregulated in subcluster 1, and amongst them, fourteen have been associated with tumorigenesis (Fig 6E). Amongst these, five are immediate early response genes (*Ier2, Fos, Junb, Jun* and *Egr1*) that can be activated by the MAPK pathway. This pathway was also activated in our *p27^-/-^* bulk RNAseq dataset (Fig 1D). In embryonic pituitary progenitors, activation of the MAPK pathway is associated with expansion of this compartment (Haston et al., 2017). We thus performed immunohistochemistry for phosphorylated ERK and observe a clear and specific upregulation of the signal in *p27^-/-^* SCs, in contrast with both control and *Sox9^iresCreERT2/+^;Sox2^fl/+^;p27^-/-^* samples (Fig 6F). This suggests that SOX2 positively regulates activity of the pathway in SCs, either directly or indirectly, and this correlates with tumourigenesis.

Subclustering of the melanotrophs, which were more numerous than the SCs, was better associated with the different genotypes (Fig 6G, H). In agreement with the absence of tumour formation, there was a reduction in melanotroph numbers in *Sox9^iresCreERT2/+^;Sox2^fl/+^;p27^-/-^* compared to *Sox9^iresCreERT2/+^;p27^-/-^* but these were still more numerous than in wild-type, in agreement with the hyperplasia observed in compound mutants (Fig 6G). Furthermore, while melanotrophs are genetically equivalent in *Sox9^iresCreERT2/+^;p27^-/-^* and *Sox9^iresCreERT2/+^;Sox2^fl/+^;p27^-/-^* samples, their transcriptome is clearly different, further confirming the SOX2-dependent effect of SCs on melanotrophs. There were 624 DEGS in cluster 0 (*Sox9^iresCreERT2/+^;p27^-/-^*) melanotrophs compared to clusters 2 (*Sox9^iresCreERT2/+^*) and 1 (*Sox9^iresCreERT2/+^;Sox2^fl/+^;p27^-/-^*) (Fig 6I). Pathway analysis of the 593 downregulated genes in *Sox9^iresCreERT2/+^;p27^-/-^* melanotrophs revealed that vesicular and endosomal traffic, and cytoskeleton remodelling, as we observed in our bulk analysis (Fig 1D, 2D), were predominantly affected, suggesting that the differentiated, secretory phenotype of melanotrophs was restored upon reduction of SOX2 in SCs (Fig 6J, upper). In agreement with this hypothesis, *Tbx19* was amongst the downregulated genes in *Sox9^iresCreERT2/+^;p27^-/-^* melanotrophs. Recovery of normal expression levels of *Tbx19* upon *Sox2* downregulation in SCs was further validated by quantitative PCR (Fig 6K). Finally, pairwise comparison between cluster 0 and 1, comprising *Sox9^iresCreERT2/+^;p27^-/-^* and *Sox9^iresCreERT2/+^;Sox2^fl/+^;p27^-/-^* cells respectively, revealed a significant enrichment in process networks associated with translation and transcription (Fig 6J, bottom), suggesting reduced levels of both transcription and translation in *p27^-/-^* cells, restored in melanotrophs upon reduction of *Sox2* levels in SCs. SCs and CSCs are known to have reduced levels of translation. Moreover, SOX2 was recently shown to be directly involved in alterations in the rate of protein synthesis in a mouse model of squamous tumour initiating cells (Sendoel et al., 2017). This suggests that SOX2 may play a similar role in *p27^-/-^* melanotrophs, acting as a repressor of translation and transcription of genes related to differentiation.

Altogether these data show that SOX2 upregulation in *p27^-/-^* SCs results in the induction of a dedifferentiated, pro-tumoral phenotype in melanotrophs. In SCs, SOX2 overexpression leads to increased MAPK pathway activation, which correlates with and may mediate the role of SOX2 in tumorigenesis, at least in *p27^-/-^* IL, but maybe also in other contexts.

## Discussion

The regulation of *Sox2* expression is extremely complex (Kondoh and Lovell-Badge, 2015) and variation of its expression levels have contrasting consequences in cancers (Wuebben and Rizzino, 2017). We previously uncovered that P27 drives *Sox2* repression and that deletion of one copy of *Sox2* in *p27^-/-^* animals rescues deleterious aspects of the *p27^-/-^* phenotype, in particular tumours of the pituitary, which are the most frequent lesions in this model (Li et al., 2012). Here we have further dissected the role of SOX2, and also demonstrated the importance of *Srr2* for the SOX2-P27 interaction. We have shown that SOX2 is required independently both in melanotrophs and SCs for IL tumours to develop in *p27^-/-^* animals (see model, Fig. 7). The requirement for SOX2 in SCs is particularly interesting, because it implies that SCs, upon loss of P27, act on melanotrophs to promote tumour formation. To better analyse this aspect, we performed a single cell transcriptomic analysis to simultaneously explore the SOX2-dependent tumour inducing effect within SCs, and the consequences for melanotrophs. These analyses revealed that in *p27^-/-^* SCs, SOX2 upregulation leads to MAPK pathway overactivation. In this model, increased proliferation and alteration of melanotroph cell identity are clearly mediated by the SCs, and this correlates with tumour formation. The pleiotropic, pro-tumorigenic effects of SOX2 in distinct cell types in the absence of P27 demonstrate that it is a crucial mediator of tumourigenesis and hence, as suggested before, a good target for anti-tumoral treatments. However, we also show that SOX2 only appears to be a relevant target in a subset of tissues when P27 is lost or its expression decreased, such as the pituitary and duodenum. It would be interesting to examine whether SCs are able to induce tumour development in the others organs comprised in this subset, as they do in the pituitary. Understanding the reasons behind the tissue specificity for the SOX2-P27 interaction could help to define pre-existing factors favoring tumorigenesis. Our results suggest that derepression of *Sox2* in the absence of P27 leads to IL tumours. However this happens only in some cell types, while expression of *p27* is ubiquitous, likely illustrating the multiple anti-proliferative roles of P27 that are distinct from *Sox2* repression (Fabris et al., 2015)(Nguyen et al., 2006). Differential expression of P27 interactors and/or targets, and of other cell cycle repressors may explain tissue-specific features of the *p27^-/-^* phenotype.

**Figure 7.**
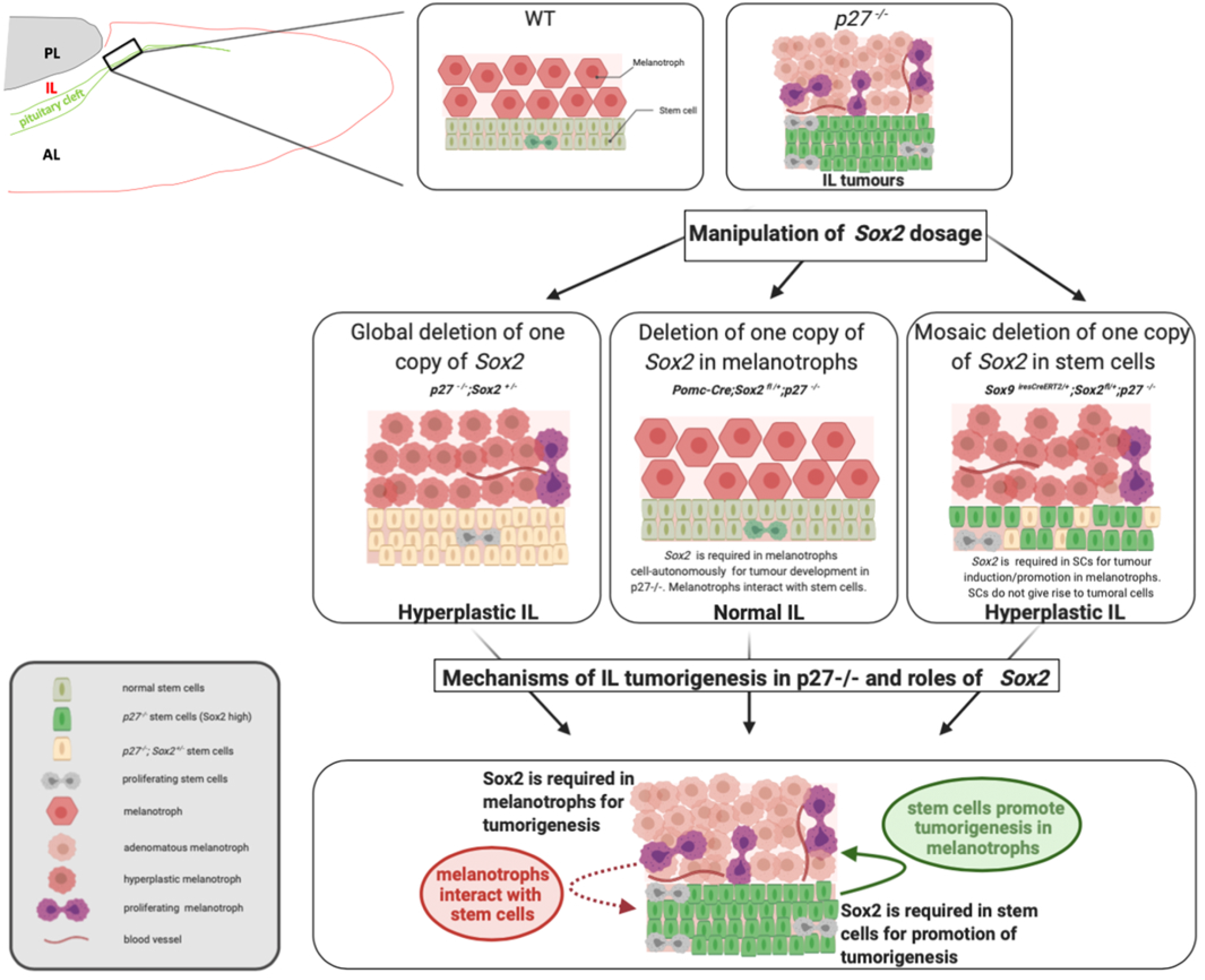
Model recapitulating the consequences of *Sox2* dosage modulation during IL *p27^-/-^* tumorigenesis and its proposed roles. In normal IL, SOX2 is expressed at high levels in SCs, that proliferate rarely, and at low levels in melanotrophs, that do not self-renew {Langlais, 2013 #482}. In *p27^-/-^* IL, melanotroph tumors develop, with ectopic vascularization; increased SCs proliferation is also observed. SOX2 levels are increased in both cell types, while expression of the differentiation markers TBX19 and POMC is reduced in melanotrophs. Removal of one copy of *Sox2* in *p27^-/-^* animals (*p27^-/-^;Sox2^+/-^*) results in impairemnt of tumorigenesis, while hyperplasia is still observed. Conditional deletion of *Sox2* in melanotrophs (*Pomc-Cre; Sox2^fl/+^;p27^-/-^*) prevents hyperplasia and tumor formation, therefore SOX2 is required cell-autonomously in *p27^-/-^* melantrophs for tumor formation (Fig. 4). Furthermore, modulation of SOX2 levels in SCs in this model suggests that melanotrophs interact with SCs (dotted arrow in proposed model). Conditional deletion of *Sox2* in SCs (*Sox9^iresCreERT2/+^;Sox2^fl/+^;p27^-/-^*) also impairs tumorigenesis, reduction of *Tbx19* levels, and SCs overproliferation but hyperplasia is still observed (Fig. 5, 6). This shows that SCs promote tumor formation and loss of differentiated features, while not giving rise to tumorigenic cells themselves, as our lineage tracing experiments show. This tumor-inducing role of SCs (arrow in proposed model) depends on SOX2. Therefore SOX2 is required both in melanotrophs and SCs for respectively cell-autonomous and tumor promoting functions following loss of P27. This model was created with BioRender.com.

In pituitary endocrine cells, SOX2 expression appears to be a prerequisite for tumoral development in absence of *p27*, since melanotrophs are the only such cell type where SOX2 is detectable (Goldsmith et al., 2016). While melanotrophs are also uniquely characterized by their non-proliferative status (Langlais et al., 2013), levels of P27 are unexpectedly relatively low in melanotrophs. This suggests that in *p27^-/-^* mutants, an active *Sox2* locus normally under P27 control, coupled with a relaxed control of the cell-cycle renders melanotrophs particularly prone to tumorigenesis. In agreement, independent deletion of genes encoding for different negative cell cycle regulators show that IL is particularly sensitive to tumoral development (Quereda and Malumbres, 2009). Furthermore derepression of *Sox2* is also observed in *pRb^+/-^* pituitary tumors (Vilas et al., 2015). It would be interesting to examine whether SOX2 is required for IL tumorigenesis in other cell cycle regulator mutants (Quereda and Malumbres, 2009). Conversely we can now screen for cell types satisfying the relevant criteria, including low cell-turnover, low negative cell cycle regulator levels, and presence of SOX2, and examine for tumorigenic potential.

To further characterise the molecular mechanisms involved in the SOX2-P27 interaction, we have deleted *Srr2*. SOX2 itself, c-myc and members of the POU transcription factor family bind to *Srr2* to promote expression of the gene (Miyagi et al., 2004). Beside P27 (Li et al., 2012), other negative regulators of *Srr2* have been characterised such as P21 in neural SCs (Marques-Torrejon et al., 2013) and pRb in the pituitary (Vilas et al., 2015). Analysis of *p27^-/-^; Srr2^del/del^* animals show that the repressive action of P27 is mediated by *Srr2* and in absence of *p27*, Srr2 activators drive *Sox2* expression; this then leads to pituitary tumours and retina defects (see model, Fig 3H). Notably, loss of *Srr2* affects cell proliferation exclusively in *p27^-/-^* SCs, suggesting that the role of the enhancer is more important in SCs, in agreement with a role for these in promoting the melanotroph tumours. Further analysis of these animals is required to assess the tissue-specific roles of *Srr2*, particularly in neural SCs where this enhancer is active (Miyagi et al., 2006),(Marques-Torrejon et al., 2013) (Domingo-Muelas A., M-A V. et al, *in preparation*).

Our study highlights common and cell-specific roles of SOX2 during tumorigenesis. Both in *p27^-/-^* melanotrophs and SCs, SOX2 upregulation is associated with overproliferation. The association of SOX2 with dedifferentiation and overproliferation has been observed in many tumours (Wuebben and Rizzino, 2017), as we also observe here in melanotrophs. In pituitary SCs, our results suggest that this overproliferation is linked to MAPK overactivation, something that has been shown in embryonic pituitaries expressing an activated form of *Braf* (Haston et al., 2017). How SOX2 derepression induces MAPK activation is less clear; MAPK pathway activation has been linked to induction of *Sox2* expression (Wang et al., 2018), but the molecular mechanisms undelying the converse have not been characterized. While SOX2 has been associated with CSC properties in some other tumours (Schaefer and Lengerke, 2020), we report here its requirement in SCs for these to promote tumorigenesis in neighbouring melanotrophs. Manipulation of MAPK in SC in *p27^-/-^* animals is now required to characterise their effect on SCs and melanotroph transformation.

In parallel with the partial melanotroph dedifferentiation observed here, we previously reported a very clear dedifferentiation of melanotrophs and corticotrophs following cell autonomous NOTCH pathway activation (Cheung et al., 2018a). While NOTCH is unlikely to be involved here, it is remarkable that we also observed increased expression of SOX2 in dedifferentiating melanotrophs. The POMC lineage was the only pituitary lineage to display such plasticity, suggesting that in addition to being more sensitive to situations where proliferative control is unlocked, melanotroph fate is also relatively unstable. The importance of this SOX2-dependent plasticity was recently demonstrated in prostate cancer where cell-autonomous derepression of SOX2 induces a cell fate change underlying chemoresistance (Ku et al., 2017),(Mu et al., 2017).

In conclusion, our study reveals how *Sox2* derepression, independently in two adjacent cell types, underlies tumorigenesis in one of them. Moreover, our work reveals the existence of reciprocal interactions between melanotrophs and flanking SCs to orchestrate tumour development. This highlights the complexity of the mechanisms triggering tumorigenesis and the central role of SOX2 in this process. Furthermore pituitary SCs, which are not tumoral, but have a tumour inducing activity, represent good anti-tumoural treatment targets, highlighting the importance of detailed characterisation of mechanisms underlying tumour formation to decipher possible anti-tumour strategies. While we have focused our attention on the IL, where we have had the tools to explore the mechanisms leading to tumour formation, SOX2 activity is often associated with tumours in other tissues and it would be interesting to explore whether its role and the mechanisms are similar.

## Acknowledgements

We are grateful for help and support from all members of the Lovell-Badge’s lab. We thank Jacques Drouin (Institut de Recherches Cliniques de Montreal) for sharing the Pomc-Cre animals and the pituitary *Pomc* regulatory element; Francis Poulat (Institut de Génétique Humaine, Montpellier) for the generous gift of SOX9 antibody; and Donald Bell for his expertise on image quantification. We are grateful to Dr. A.F. Parlow and the NHPP for reagents for the sandwich ELISA and immunofluorescence experiments. We also thank Biological Services, Flow cytometry, Advanced Sequencing, and Advanced light microscopy platforms at the Francis Crick Institute, for their excellent assistance and technical support. This work was supported by the Medical Research Council, U.K. (U117512772, U117562207 and U117570590), the Francis Crick Institute which receives its core funding from Cancer Research UK (FC001107), the UK Medical Research Council (FC001107), and the Wellcome Trust (FC001107), and the Worldwide Cancer Research charity (grant 13-1270).

## Material and Methods

### Ethics Statement

All experiments carried out on mice were approved under the UK Animal (scientific procedures) Act (Project licence 80/2405 and 70/8560).

### Mice

*Cdkn1b^tm1Mlf^* (*p27^-^*) (Fero et al., 1996), *Sox9^tm1(Cre/ERT2)Haak^* (*Sox9^iresCreERT2^*) (Furuyama et al., 2010), *Tg*(*PomC-Cre*)*Dro*(*Pomc-Cre*) (Langlais et al., 2013), *Sox2^tm1Lpev^*(Sox2^GFP^), *Sox2^tm2Lpev^*(Sox2^fl^) (Taranova et al., 2006) and *Gt*(*ROSA*)*26Sor^tm1(EYEP)Cos^* (*Rosa26^ReYEP^*) (Srinivas et al., 2001) were maintained on mixed background. Experiments were either performed using littermates or animals with comparable backgrounds.

Cre activity was induced by tamoxifen administration for 3 consecutive days at 5mg/25g body weight /day. Pituitaries were harvested after tamoxifen treatment, as indicated in each experiment. Genotyping was performed by Transnetyx®.

### Generation of Pomc-CreERT2 transgenic mice and and *Sox2^tm1(Guide1/1:SRR2)RLB^* mice

*Tg(Pomc-cre/ERT2)Rlb* (*Pomc-CreERT2*) mice were generated by inserting a 543bp (−480/+63) region of the *Pomc* promoter (Langlais et al., 2013) upstream of the sequence coding for iCreERT2. *Pomc-CreERT2* animals were obtained by standard pronuclear injection. Tissue-specific activity of Pomc-CreERT2 was confirmed by crossing transgene positive animals with *Rosa26^ReYEP/ReYEP^* mice; one founder was chosen to establish the strain.

The *Sox2^tm1(Guide1/1:SRR2)RLB^* allele was generated using the CRISPR/Cas9 technology. Single guide RNAs (sgRNAs) were designed (www.crispr.mit.edu) to induce deletion of the mouse SRR2 enhancer (81 bp), situated 4kb downstream of *Sox2* (UCSC mm9 genome chr13: 34,552,926 - 34,553,007). Two pairs of guides were designed: pair 1(G1_5’+G1_3’) and pair 2 (G4_5’+G5_3’) (Table S2) and independently injected into zygotes with Cas9 mRNA as previously described (Gonen et al., 2018) to produce two different mouse strains with the same *Srr2* deletion, in order to discriminate between the effects of the desired mutation, which would be common to both guides, and potential off-target effects of the guides which would be in contrast specific to a particular pair. Founders were genotyped for SRR2 deletion by PCR and mosaicism estimated using the MiSeq system (Illumina) (Table S3, adaptor sequence in bold). Two founders were chosen to establish two independent strains.

### Immunofluorescence, EdU staining, image acquisition and pre-processing

Immunofluorescence was performed as previously described (Rizzoti et al., 2004). Mice were perfused with 4% w/v paraformaldehyde in phosphate-buffered saline (PBS), pituitaries harvested and cryosectioned at 12 *μ*m. Sections were blocked with blocking solution (10% v/v donkey serum in PBS/0.1% v/v Triton X-100; PBST) for 1 hr, then incubated with primary antibodies in 10% blocking solution overnight at 4°C. Primary antibodies were used at the following dilutions: rat anti-GFP (Nacalai Tesque) at 1:1000; rabbit anti-POMC (NHPP) at 1:500; mouse anti-POMC (Barratt et al.) 1:500, Mouse anti-PAX7 (DSHB) 1:100, goat anti-SOX2 (Immune System) at 1:300, rabbit anti-SOX9 (gift from F. Poulat, Institut de Génétique Humaine, Montpellier) at 1:300, rat anti-CD31 (Furlan et al.) 1:50, p27kip C19 (SantaCruz) 1:300, rabbit anti-ki67 (Abcam) 1:1000. Sections were washed in PBST then incubated for 1 hr at room temperature with the corresponding anti-rat, anti-goat or anti-rabbit secondary antibody conjugated to Alexa-Fluor 488, 555, 594 or 647 in 10% blocking solution with 1 *μ*M 4’, 6-diamino-2-phenylindole (DAPI).

Cell proliferation was analysed following a one-hour EdU pulse in E18.5 embryos (30*μ*g/g body weight), or two intraperitoneal daily injections for 3 consecutive days in adult. Adult pituitaries were harvested at day 4. EdU incorporation was detected using the Click-iT EdU imaging kit following the manufacturer instructions (Thermo Fisher Scientific). Sections were washed with PBST and mounted using Aqua-Poly/Mount (Polysciences, Inc., Warrington, PA, USA).

Images were acquired using a Leica SPE confocal microscope or Olympus VS120 Slide Scanner. Settings were established during the initial acquisition. All images taken from the Leica SPE confocal microscope were pre-processed using ImageJ (maximum z-projection) and the ones taken from VS120 Slide Scanner were processed using Qu-path.

### Immunohistochemistry

Pituitary glands were fixed in 10% buffered formaldehyde for 16 hours and embedded in paraffin. For staining, 4μm sections were de-paraffinized using xylene and rehydrated through a graded series of ethanol. Antigen retrieval was performed for 20 min at high temperature in either 0.01M citrate buffer (pH6) or Tris-EDTA (10mM Tris base, 1mM EDTA solution, pH9), depending on the antibody. The following antibodies were used: Sox2 (AF2018, R&D), 1:100, on sections treated with antigen retrieval buffer (Ventana) for 48 minutes, followed by a one hour primary antibody incubation. IHC was performed on the Discovery Ultra Ventana platform (Roche). P-ERK1/2 (4370, Cell Signalling Technology) 1:100 O/N at 4°C, antigen retrieval was performed in the microwave for 23 minutes with 0.01M citrate buffer pH6. Goat anti-rabbit secondary antibody (BA-1000, Vector) was incubated 1:250 for 45 minutes at RT and then signal amplification and HRP detection were performed using the ABC kit (Vectorlabs) for 30 minutes at RT. This IHC was performed manually. Samples were blocked using 1% BSA and incubated overnight at 4°C with the desired antibody, or in blocking buffer for controls. Finally, slides were incubated with the secondary antibody for one hour and washed three times with PBS. For colorimetric staining with diaminobenzidine (DAB) slides were incubated with peroxidase substrate and mounted.

### Cell Countings

For proliferation assays quantification, three different fields were chosen on different sections (sections always include IL, SCs flanking the cleft and AL). Within these, the numbers of SOX2-positive, SOX9-positive, EdU-positive, and SOX2;EdU;SOX9 triple-positive cells, representing proliferative stem cells, or SOX9 negative, SOX2;EdU double-positive cells, representing proliferative melanotrophs, were counted blindly and manually.

In EdU injected *Pomc-CreERT2; Sox2^fl/+^;p27^-/-^;Rosa26^ReYEP/+^* mutants, a total of around 1000 PAX7 positive cells was counted from fields randomly chosen and encompassing the whole IL. The number of EdU;PAX7-double-positive or EdU;PAX7;eYFP-triple-positive cells was counted blindly and manually, and the IL surface area was determined using the Fiji software.

The Fiji software was used to pseudocolor the different channels in unprocessed micrographs. The Qupath software (Bankhead et al., 2017) was also used for image analysis. Briefly, areas of interest were defined after POMC immunostaining for the intermediate lobe, or SOX2;SOX9 for the SC layer. Within the areas of interest, the centre of cell nuclei was then identified as maxima in a filtered DAPI image. Nuclear boundaries were assigned by a propagation algorithm, and then expanded by ~1 micron to define sampling areas. The following data were then recorded: (i) average pixel intensities for each data channel over each sampling area, representing one cell, (ii) the size of the sampling areas; and (iii) the number of positive cells for each data channel over each sampling area.

### Pituitary Dissociation for FACS, sc-RNAseq and Sphere Assay

Pituitaries were harvested, the posterior lobe was removed and the anterior and intermediate lobes isolated and incubated separately in a solution of papain (10108014001, Sigma-Aldrich, 1mg/ml in HBSS) for 15 min at 37°C in presence of 10□g/ml of both DNAse (10104159001, Thermo Scientific) and Rock Inhibitor (M1817, Abmole Bioscience). The papain solution was then removed, and mechanical dissociation was performed on ice in pituisphere medium (Fauquier et al., 2008). Pituispheres were derived as previously described (Fauquier et al., 2008) and spheres counted manually and blindly after 1 week in culture.

### mRNA extraction and reverse transcriptase-PCR (rt-qPCR)

Total mRNA was extracted from dissected IL using the RNeasy Micro kit (Qiagen) according to the manufacturer’s protocol. The extracted RNA was reverse-transcribed into cDNA using the Superscript VILO cDNA synthesis kit (Thermo Fisher Scientific, Figure 2) or SMART-Seq v4 Ultra Low Input RNA Kit (Takara Bio USA, Figure 5) according to the manufacturer’s protocol.

### Quantitative real-time PCR (RT-qPCR)

Each sample was assayed in technical duplicate with each tube containing diluted template cDNA, 250 nM primers and 1xAbsoluteSybrGreen ROX mix (Thermo Fisher Scientific). Each sample was assayed for the genes of interest together with the reference housekeeping gene *Gapdh* (Table S4). Relative expression of the genes of interest was calculated by normalisation of the detected expression value to the geometric mean of the reference genes using the ΔΔCt method (Livak and Schmittgen, 2001). Data is shown as mean±SEM with the number of biological samples indicated in each figure. Sidak’s (Figure 1I) and Tukey’s multiple comparison tests (Figure 2H and 5K) were used to assess significance of the data.

### Radioimmunoassay (RIA)

Pituitaries were homogenized in phosphate-buffered saline and hormonal contents measured by RIA (McGuinness et al., 2003) using National Hormone and Pituitary Program reagents kindly provided by A.L. Parlow for ACTH and GH and using alpha MSH RIA kit (RB303-Invitech).

### In-situ hybridization (RNAscope)

Fluorescent in situ hybridisations were performed manually using RNAscope (Multiplex Fluorescent Reagent Kit v2, 320293) on cryosections according to the manufacturer’s protocol. Images were analysed and quantified using the Fiji package.

### Statistical analyses

Statistical analyses were performed using Prism v.8.0c (GraphPad Software, USA). To examine significance of the data, tests were selected according to the experiment analysed. For survival curves generated using the Kaplan Meier method, log-rank tests were applied (Fig. 2B). When comparing two groups of values with normal distribution, unpaired t-test was performed (Figure; 1H, 3H, 3I and in Supplementary figure; 1B & 4B). Angular transformation was applied to compare percentages followed by unpaired t-test (Figure; 1L, 1M, 2E and 3D. When comparing two groups of values where the distribution was non-parametric, Mann Whitney test was performed (Figure; 1C (upper graph), 1G, 1J). For multiple comparisons, analysis of variance (ANOVA) was implemented; Sidak’s test was performed when comparing two groups (genotype) and differences between compartments (AL, SC cleft or IL) in Figure; 1C (bottom panel), 1K, 1O. To analyse larger groups, Tukey’s multiple comparison test was implemented (Figure 2G, 2M & O, 3C, 3K, 4C, 4D, 4E and Supplementary Figure; 2B, 4A). All results are represented as means ± standard deviation (SD) for raw data and means ± standard error of the mean (SEM) for graphs. Standard significance levels were used: *p<0.05, **p<0.01, ***p<0.001, ****p<0.0001.

### RNA sequencing sample preparation

#### Bulk RNA sequencing

RNA was extracted from dissected male and female IL or FAC sorted cells using the RNeasy Micro kit (Qiagen) according to the manufacturer’s protocol. RNA quality was assessed using the Agilent RNA 6000 Pico Kit (Agilent Technologies). cDNA was generated using Ovation RNA-seq System V2 (Tecan, 7102-A01), libraries were constructed using Ovation Ultralow System V2 (Tecan, 0344NB-A01) according to the manufacturer’s instructions. Libraries were quantified using the TapeStation (Agilent) and pooled in equimolar proportions. Library were sequenced on an Hiseq4000 (Illumina), to achieve an average of 25 million reads per sample.

#### Single-cell RNA sequencing

Male and female IL cells were dissociated as described above. Libraries were generated using Chromium Single Cell 3’ kit v3 (10x genomics, 100092) according to the manufacturer’s instructions. Both cDNA and libraries were quantified using the TapeStation (Agilent) and sequenced on an Hiseq4000 (Illumina), to achieve an average of 50,000 reads per cell.

### Bioinformatics analysis

#### Bulk RNA sequencing

The sequencing was performed on biological duplicates or triplicates for each data point. The RSEM package (version 1.3.30) (Li and Dewey, 2011) was used in conjunction with the STAR alignment algorithm (version 2.5.2a) (Dobin et al., 2013) for the mapping and subsequent gene-level counting of the sequenced reads with respect to Ensembl mouse GRCm.38.89 version transcriptome. All parameters for RSEM were run as default except “-forward-prob” which was set to 0.5. Normalisation of raw count data and differential expression analysis was performed with the DESeq2 package (version 1.18.1) (Love et al., 2014) within the R programming environment (version 3.4.3) (Team, 2008). Differentially expressed genes were defined as those showing statistically significant differences between pairwise groups if the adjusted P value was less than 0.05 (FDR < 0.05). Differentially expressed genes were taken forward and their pathway and process enrichments were analysed using Metacore (https://portal.genego.com). A hypergeometric test was used to determine statistical enriched pathways and processes and the associated P-value was corrected using the Benjamini–Hochberg method.

#### Single-cell RNA sequencing

10x CellRanger (version.3.0.2) was used to generate single cell count data for each genotype (*Sox9^iCreERT2/+^;Rosa26^ReYEP/+^, Sox9^iCreERT2/+^;p27^-/-^ Rosa26^ReYEP/+^* and *Sox9^iCreERT2/+^;Sox2^fl/+^;p27^-/-^ Rosa26^ReYEP/+^*) using a transcriptome built from the Ensembl mouse GRCm38 release 89. All subsequent analyses were performed in R v.3.6.0 using the Seurat (v3) package (Stuart et al., 2019). Primary filtering was performed on each dataset by removing from consideration: cells expressing fewer than 50 genes and cells for which mitochondrial genes made up greater than 10% of all expressed genes. Each dataset was normalised using the’ LogNormalize’ function, with a scale factor of 10,000. The top 2000 highly variable genes were found using the ‘FindVariableGenes’ function and the data centred and scaled using the ‘ScaleData’ function. PCA decomposition was performed and after consideration of the eigenvalue ‘elbow-plots, the first 20 components were used to construct Uniform Manifold Approximation Projection (UMAP) plots. Clusters relating to Melanotrophs and Stem Cells were identified, using the expression of known markers [melanotroph markers: Pomc, Pax7 and Pcsk2; stem cell markers: Sox2 and Sox9], and these clusters were integrated across the three genotypes (*Sox9^iCreERT2/+^;Rosa26^ReYEP/+^, Sox9^iCreERT2/+^; p27^-/-^;Rosa26^ReYEP/+^* and *Sox9^iCreERT2/+^; Sox2^fl/+^; p27^-/-^ Rosa26^ReYEP/+^*) using Seurat 3’s standard integration workflow. Differentially expressed genes between clusters across genotypes were determined using the ‘FindMarkers” function.

### Data availability

The RNA-sequencing datasets have been deposited in the Gene Expression Omnibus with the grouped accession number GSE152010. The bulk RNAseq datasets are GSE152007 (*p27^-/-^* IL compared to wild-type, two- and seven-month old) and GSE152008 (*p27^-/-^* IL compared to *p27^-/-^; Sox2^+/^* and wild-type, two-to three-month old); the single-cell RNA-sequencing datasets have been deposited with accession number GSE152009.

---

**Supplementary figure 1:**
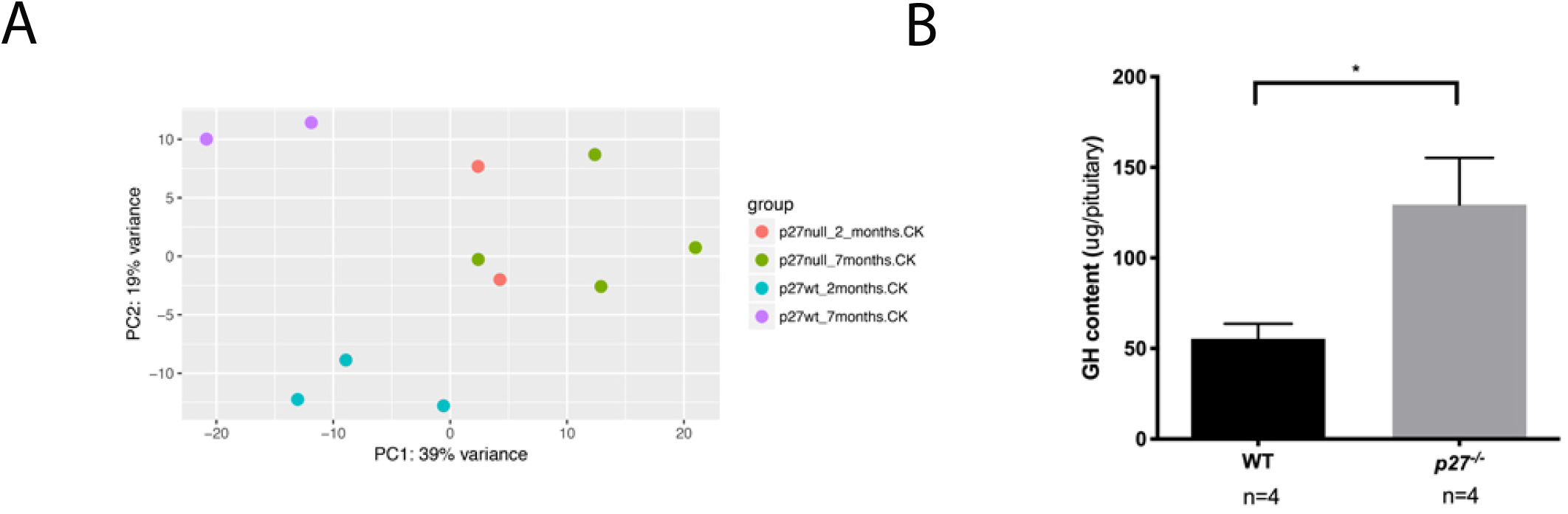
Analysis of *p27^-/-^* pituitaries. **A)** Principal Component Analysis (PCA) plots of bulk RNA-seq data of 2- and 7-month old wildtype and *p27^-/-^* IL samples. There is a clear segregation of samples according to genotype, even before tumour formation (2 month-old). Furthermore while wild-type samples segregate according to age, *p27^-/-^* samples appear more similar, independently of the age of the animal. **B)** GH content were measured by RIA in 2-month old male pituitaries. *p27^-/-^* pituitary glands contain higher GH levels (*p=0.0340, n=4 in each group) which is consistent with gigantism affecting these mutants.

**Supplementary figure 2:**
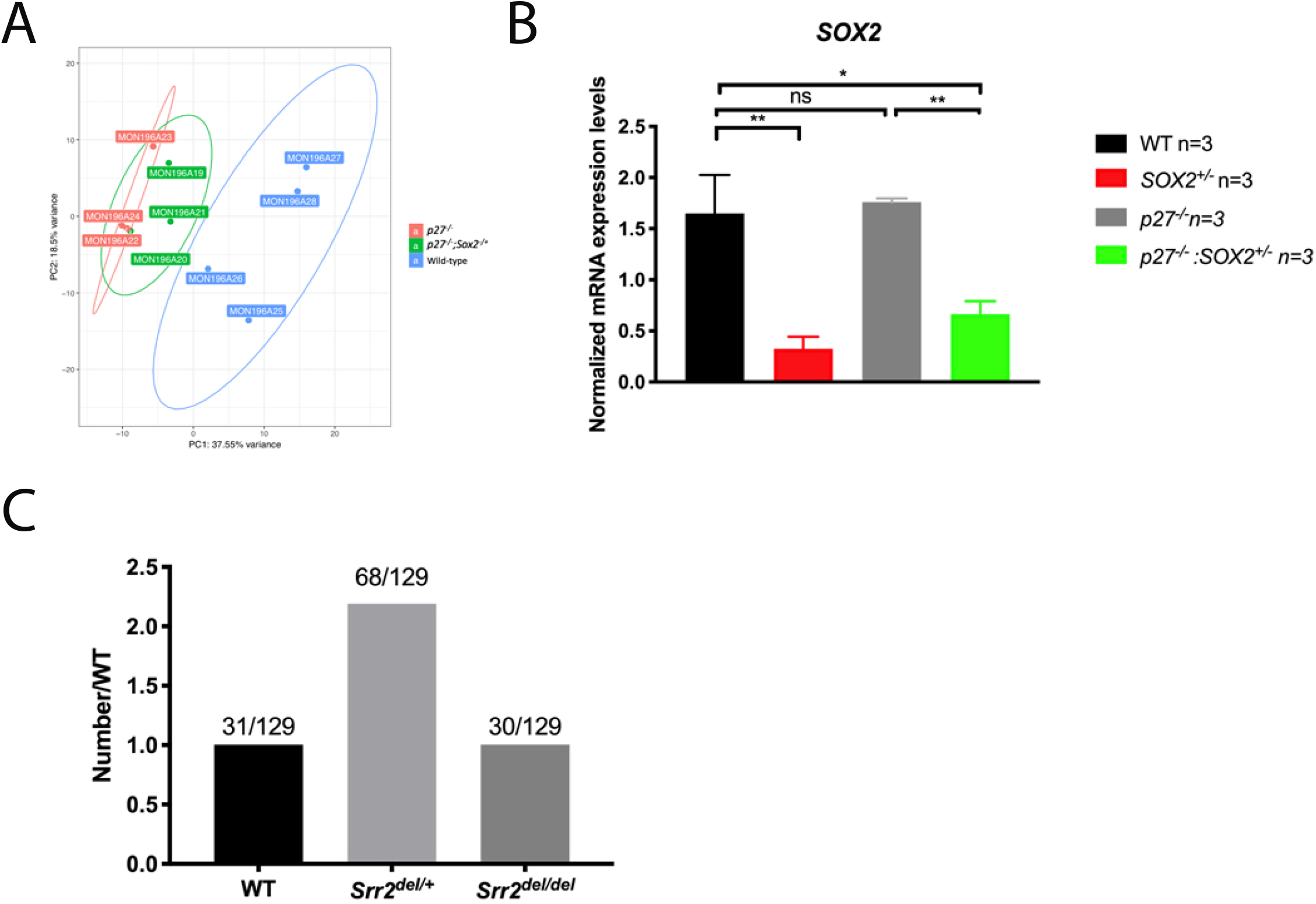
Analysis of *p27^-/-^;Sox2^+/-^* and *p27^-/-^;Srr2^del/del^* pituitaries. **A)** PCA plots of bulk RNA-seq data of 2-to 3-month old wild-type, *p27^-/-^* and *p27^-/-^; Sox2*^+/-^ ILs. *p27^-/-^; Sox2*^+/-^ samples are clustering between wild-type and *p27^-/-^* ones. **B)** RT-qPCR analysis of *Sox2* in wild-type, *Sox2^+/-^, p27^-^*, and *p27^-/-^; Sox2*^+/-^ in 2-to 3-month old ILs. *Sox2* expression levels decrease in *Sox2^+/-^* and *Sox2^+/-^;p27^-/-^* vs w*t* or *p27^-/-^*. We did not observe an increase in *Sox2* levels in *p27^-/-^* using this technique. This may be because IL comprises two cell types expressing *Sox2* at different levels: a high proportion of melanotrophs, expressing it at low levels, and SCs, expressing it at high levels. In *p27^-/-^* animals there are many more melanotrophs than in wild-type, but these, despite *Sox2* derepression, are still expressing *Sox2* at lower levels than SCs (Fig. 1A). The difference in proportion in both cell types between wildtype and mutant may explain the inability to observe *Sox2* derepression. **C)** Genotypes of offspring obtained from *Srr2^del/+^* heterozygous (XX) and (XY) intercrosses.

**Supplementary figure 3:**
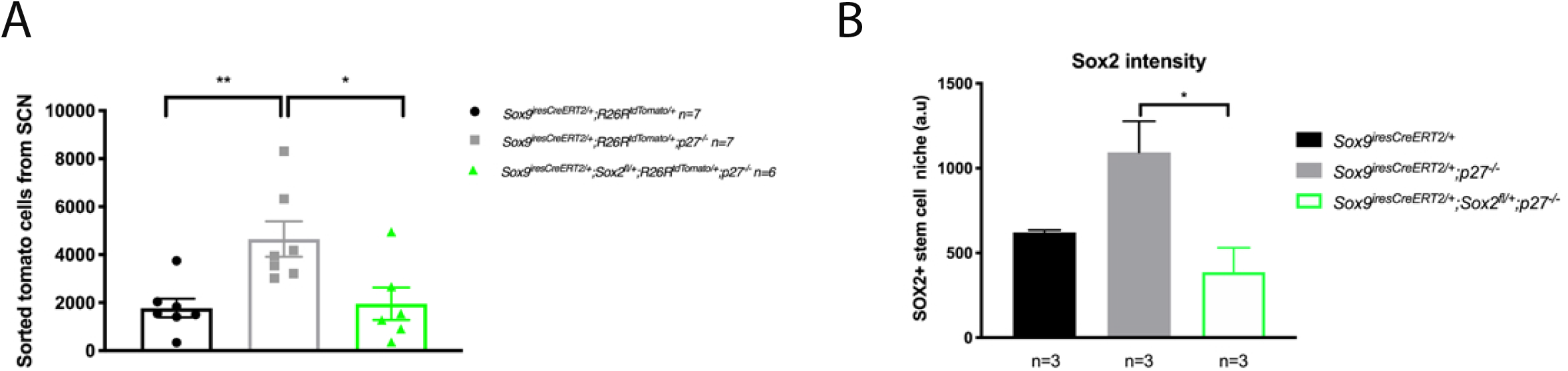
Analysis of stem cells in *Sox9^iresCreERT2/+^;Sox2^fl/+^;p27^-/-^* pituitaries. **A)** Analysis of tomato positive cells by flow cytometry in *Sox9^iCreERT2/+^;Rosa26^tdTomato/+^, Sox9^iCreERT2/+^; p27^-/-^;Rosa26^tdTomato/+^* and *Sox9^iCreERT2/+^;Sox2^fl/+^;p27^-/-^;Rosa26^tdTomato/+^* dissected IL from 3.5-to 5-month old animals that were treated by tamoxifen at 1month-old. The number of tomato-positive FACSorted stem cells is reduced in *Sox9^iCreERT2/+^;Sox2^fl/+^;p27^-/-^;Rosa26^tdTomato/+^* (1953±1658, n=6) compared to *Sox9^iCreERT2/+^;p27^-/-^;Rosa26^tdTomato/+^* samples (4648±1954, n=7), to values comparable to control. (**p=0.0094 and *p=0.0189). **B)** Quantification of SOX2 immunofluorescence intensity in the stem cell layer in *Sox9^iCreERT2/+^*(621±23 a.u, number of cells counted 1214, n=3 animals), *Sox9^iCreERT2/+^:p27^-/-^* (1093±317 a.u, number of cells counted 899, n=3) and *Sox9^iCreERT2/+^;Sox2^fl/+^;p27^-/-^* (387±246 a.u, number of cells counted 1265, n=3) pituitaries of 7 to 12 month-old animals. *Sox9^iCreERT2/+^;Sox2^fl/+^;p27^-/-^* express lower levels of Sox2 (*p=0.02).

**Supplementary Table 1:** Genes differentially expressed in the SC clusters (Single-cell RNAseq analysis) Genes differentially expressed in cluster 1 (comprising mostly *Sox9^iresCreERT2/+^:p27^-/-^* cells) but unaffected in cluster 2 (comprising mostly *Sox9^iresCreERT2/+^;p27^-/-^;Sox2^fl/+^* cells) compared to cluster 0 (comprising mostly control *Sox9^iresCreERT2/+^* cells) are listed. The percentage of cells expressing the gene of interest in each cluster is represented, followed by the significance of differential expression versus control.

| Gene | WT_vs_P27N_logFC | WT_vs_S2CK_logFC | WT.percent | P27N.percent | S2CK.percent | P27N.p_val_adj | S2CK.p_val_adj |
| --- | --- | --- | --- | --- | --- | --- | --- |
| *Ier2* | -2.142136022 | -0.477223619 | 0.411 | 0.922 | 0.652 | 2.30E-15 | 1 |
| *Fos* | -2.053977157 | -0.853951493 | 0.221 | 0.828 | 0.449 | 1.56E-13 | 1 |
| *Jun* | -1.434398863 | -0.749281189 | 0.453 | 0.906 | 0.797 | 1.47E-09 | 0.312946692 |
| *Junb* | -1.460070386 | -0.778420638 | 0.263 | 0.828 | 0.594 | 3.16E-09 | 1 |
| *Egr1* | -1.54460355 | -0.561252733 | 0.116 | 0.641 | 0.362 | 4.96E-09 | 1 |
| *Id1* | -1.373226163 | -0.666162636 | 0.011 | 0.453 | 0.174 | 7.26E-08 | 1 |
| *Uba52* | -0.88893215 | 0.528001458 | 0.547 | 0.953 | 0.652 | 5.60E-07 | 1 |
| *Hes1* | -0.93186644 | -0.196595776 | 0.389 | 0.828 | 0.681 | 2.33E-05 | 1 |
| *Socs3* | -1.268278907 | -0.678321742 | 0.063 | 0.469 | 0.348 | 2.83E-05 | 0.075346417 |
| *Id3* | -1.722005492 | -1.092530454 | 0.063 | 0.469 | 0.174 | 5.24E-05 | 1 |
| *Igfbp5* | -0.969440249 | 0.411231394 | 0.526 | 0.891 | 0.493 | 5.49E-05 | 1 |
| *Rgs2* | -1.046067021 | -0.172617514 | 0.063 | 0.453 | 0.217 | 9.50E-05 | 1 |
| *Rpl37* | -0.404226685 | -0.091294239 | 0.989 | 1 | 0.986 | 0.00010622 | 1 |
| *Tmem176b* | 0.742858292 | 0.363874608 | 0.968 | 0.984 | 0.986 | 0.000111901 | 1 |
| *Rpl38* | -0.420114841 | 0.07978908 | 0.989 | 1 | 1 | 0.000139345 | 1 |
| *Fosb* | -1.122158541 | -0.541223356 | 0.053 | 0.422 | 0.29 | 0.000169659 | 0.774302347 |
| *Atf3* | -0.866519958 | -0.507652767 | 0.021 | 0.344 | 0.203 | 0.000435014 | 1 |
| *Cd9* | 0.631709463 | 0.218219067 | 0.958 | 0.984 | 0.971 | 0.000474834 | 1 |
| *Tspo* | 0.982139941 | 0.489801839 | 0.789 | 0.625 | 0.812 | 0.00049656 | 1 |
| *Gm42418* | 1.099406939 | 0.74396475 | 0.926 | 0.828 | 0.928 | 0.000787712 | 0.210648356 |
| *Btg2* | -0.996221569 | -0.522991145 | 0.105 | 0.484 | 0.406 | 0.000967703 | 0.302739885 |
| *Meg3* | -0.917927319 | -0.364847161 | 0.116 | 0.516 | 0.377 | 0.003255314 | 1 |
| *Snhg9* | -0.726306486 | 0.374232004 | 0.232 | 0.656 | 0.275 | 0.003689219 | 1 |
| *Gadd45g* | -0.929358882 | 0.105173556 | 0.105 | 0.469 | 0.174 | 0.006306571 | 1 |
| *Epha5* | -0.716366326 | -0.22562446 | 0.053 | 0.359 | 0.145 | 0.009557697 | 1 |
| *Zfp36* | -0.875737936 | -0.29783556 | 0.126 | 0.469 | 0.29 | 0.010005941 | 1 |
| *Aldh1a2* | -0.789177177 | 0.195512656 | 0.084 | 0.438 | 0.145 | 0.011401663 | 1 |
| *Polr2m* | -0.624877875 | -0.102347863 | 0.168 | 0.547 | 0.493 | 0.016073591 | 1 |
| *Cyr61* | -0.719638566 | -0.151919162 | 0.032 | 0.312 | 0.246 | 0.019807083 | 1 |
| *Dbi* | 0.565286542 | 0.263621868 | 0.979 | 0.969 | 0.986 | 0.023267759 | 1 |
| *Apoe* | -0.759876078 | 0.408797415 | 0.453 | 0.828 | 0.507 | 0.026965005 | 1 |
| *Tle4* | -0.517427321 | 0.257254319 | 0.221 | 0.641 | 0.377 | 0.027098377 | 1 |
| *Ppia* | 0.332171961 | -0.074944715 | 0.989 | 1 | 1 | 0.027495806 | 1 |
| *Rpl6* | -0.362216237 | 0.102094813 | 0.968 | 1 | 0.986 | 0.028472481 | 1 |
| *Pcdh17* | -0.722975839 | 0.090436245 | 0.095 | 0.422 | 0.232 | 0.030787738 | 1 |

**Supplementary Table 2:**
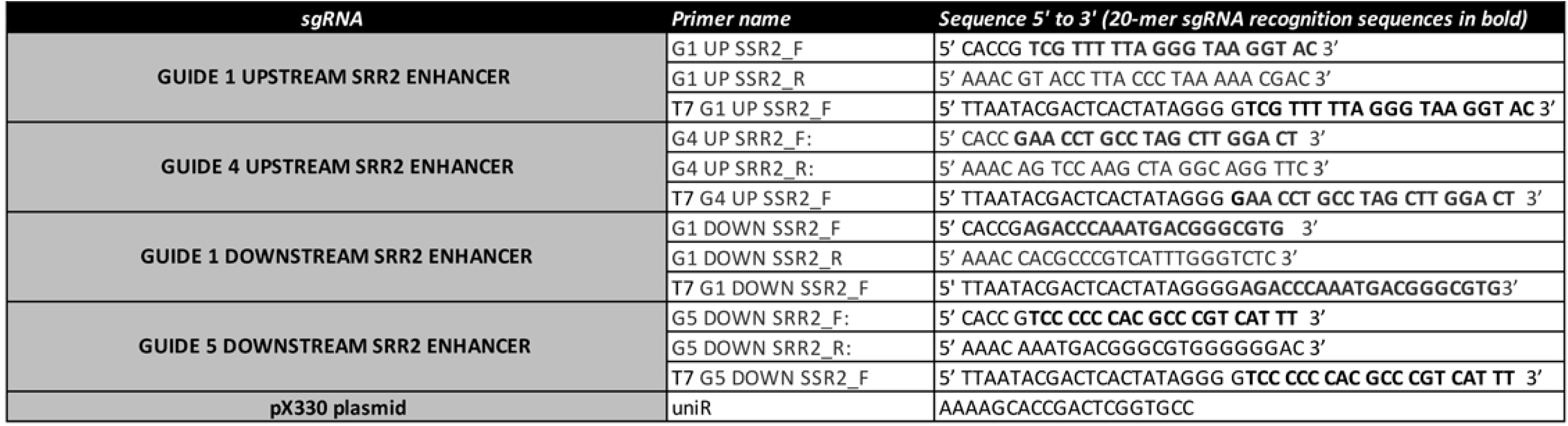
Primers used for CRISPR genome editing

**Supplementary Table 3:**
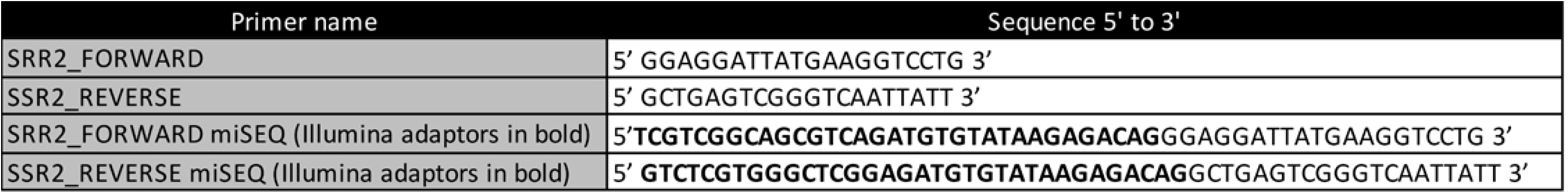
Primers used to genotype *Srr2* deleted mice.

**Supplementary Table 4:**
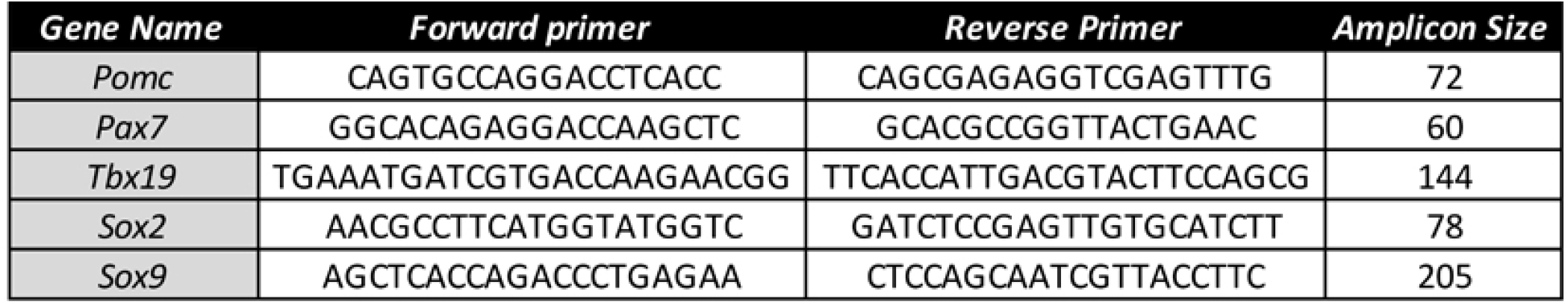
Primers used for quantitiative PCR.

